# The evolution of whole-brain turbulent dynamics during recovery from traumatic brain injury

**DOI:** 10.1101/2022.11.30.518501

**Authors:** Noelia Martínez-Molina, Anira Escrichs, Yonatan Sanz-Perl, Aleksi J. Sihvonen, Teppo Särkämö, Morten L. Kringelbach, Gustavo Deco

## Abstract

It has been previously shown that traumatic brain injury (TBI) is associated with reductions in metastability in large-scale networks in resting state fMRI. However, little is known about how TBI affects the local level of synchronization and how this evolves during the recovery trajectory. Here, we applied a novel turbulent dynamics framework to investigate the temporal evolution in whole-brain dynamics using an open access resting state fMRI dataset from a cohort of moderate-to-severe TBI patients and healthy controls (HCs). We first examined how several measures related to turbulent dynamics differ between HCs and TBI patients at 3-, 6- and 12-months post-injury. We found a significant reduction in these empirical measures after TBI, with the largest change at 6-months post-injury. Next, we built a Hopf whole-brain model with coupled oscillators and conducted *in silico* perturbations to investigate the mechanistic principles underlying the reduced turbulent dynamics found in the empirical data. A simulated attack was used to account for the effect of focal lesions. This revealed a shift to lower coupling parameters in the TBI dataset and, critically, decreased susceptibility and information encoding capability. These findings confirm the potential of the turbulent framework to characterize whole-brain dynamics after TBI and validates the use of whole-brain models to monitor longitudinal changes in the reactivity to external perturbations.

**Highlights:**

- Whole-brain turbulent dynamics capture longitudinal changes after TBI during one-year recovery period
- TBI patients show partial recovery of resting state network dynamics at large spatial scales
- Whole-brain computational models indicate less reactivity to *in silico* perturbations after TBI

## 1. Introduction

There has been a growing interest in the development and application of large-scale, non-linear brain modeling methods to resting state functional magnetic resonance imaging (fMRI) data (Breakspear, 2017; Cabral et al., 2014; Deco et al., 2018; Deco et al., 2019; Deco & Kringelbach, 2014; Deco et al., 2021; Deco et al., 2015; Kringelbach & Deco, 2020) as it offers whole-brain coverage and can be acquired in patient populations who are unable to perform cognitive tasks (Demertzi et al., 2015; Escrichs et al., 2022; López-González et al., 2021). Importantly, these methodological advancements have enabled reconceptualizing existing measures of non-linear dynamics in order to explain the complex dynamics of human brain activity. One prominent example is turbulence, which has been recently demonstrated in the healthy brain (Cruzat et al., 2022; De Filippi et al., 2021; Deco & Kringelbach, 2020) and in patients with a disorder of consciousness following brain injury (Escrichs et al., 2022). In this study, we combine a turbulent dynamics framework (model-free) with a computational modeling approach (model-based) to characterize whole-brain dynamics in moderate-to-severe traumatic brain injury (TBI) patients during a one-year recovery period.

Traumatic brain injury is associated with a wide range of cognitive deficits, such as attentional or memory impairments or executive dysfunction, that are often persistent, contribute to poor functional outcomes and have a profound impact on the quality of life of the patients (McAllister et al., 2004; Ponsford et al., 2014). The emergence of cognitive function in the brain is contingent on the integrity of the structural connectome, especially long-range white matter connections that contribute to the small-world topology underlying the spatiotemporal dynamical patterns observed in a healthy regime (Bassett & Bullmore, 2006). In addition to focal lesions, diffuse axonal injury (DAI) has been considered one of the main mechanisms of TBI (Jolly et al., 2021), leading to disturbances in long-range connections and ultimately disrupting the spatiotemporal properties of functional brain networks (Caeyenberghs et al., 2014; Hellyer et al., 2013; Jilka et al., 2014; Kinnunen et al., 2011). Such structural and functional disruption results in long-term cognitive impairment (Bonnelle et al., 2011; Jilka et al., 2014; Kinnunen et al., 2011) and may play a role in the progressive neurodegeneration associated with TBI (Sharp et al., 2014). Nevertheless, reliable biomarkers accurately predicting cognitive outcomes after TBI have not been recognized (National Academies of Sciences et al., 2022). Here, resting state fMRI-based measures like turbulence could be clinically relevant as they can be assessed repeatedly without a specific cognitive task (De Filippi et al., 2021; Deco & Kringelbach, 2020; Escrichs et al., 2022). Despite its clinical relevance, a better understanding of how TBI affects whole-brain dynamics longitudinally is still needed.

The shift towards describing the impact of brain injury in terms of changes in whole-brain dynamics holds great promise to bestow a common framework not only to characterize cognitive deficits after TBI but, more generally, to provide insights into the mechanistic principles that support brain function. One widely used approach to studying whole-brain dynamics is based on metastability, a dynamical regime where brain regions engage and disengage flexibly over time showing transient dynamical patterns (Friston, 1997; Shanahan, 2010; Tognoli & Kelso, 2014). In the context of TBI, recent evidence has shown that patients had reduced global and network-level metastability compared to controls (Hellyer et al., 2015). Critically, this reduction was associated with cognitive impairment and damage to structural connectivity suggesting that metastability may act as a conceptual bridge between brain structure and behavior (Tognoli & Kelso, 2014). However, the concept of metastability measured as the variability over time of the global level of synchronization of the brain, commonly known as the global Kuramoto order parameter of a dynamical system (Kuramoto, 1984), is limited as it cannot dissociate the dynamical patterns occurring at local spatial scales.

Recently, we have proposed an extension of metastability to account for such limitation based on turbulence theory developed first for fluid dynamics and since extended to other domains. As shown by Kuramoto (Kuramoto, 1984), coupled oscillators can be used to describe synchronization globally through metastability (Cabral et al., 2014; Deco, Kringelbach, et al., 2017) and to capture turbulence using a measure of local metastability (Kawamura et al., 2007). By combining Kuramoto’s framework with Kolmogorov’s concept of structure functions (Kolmogorov, 1941), turbulence can be calculated locally at different spatial scales in an analogy to the rotational vortices found in fluid dynamics. By taking this approach, we have found long-range correlations in higher-order task-specific regions that could be controlling a functional resting state core involving primary sensory regions (Deco & Kringelbach, 2020). Further, the results from the whole-brain computational model indicated that turbulence is a good index of the information processing capability of the brain as both reached the maximum level at the optimal fitting of the model. When applied to a population of patients with disorders of consciousness —which is often altered in severe cases of TBI— turbulence was significantly reduced at large and intermediate distances, as was the sensitivity of the brain to respond to external perturbations (Escrichs et al., 2022). Therefore, the question arises as to whether similar alterations in whole-brain dynamics could be found after TBI, across different stages of recovery.

Here, by combining empirical and computational approaches, we investigated how whole-brain dynamics are affected by TBI measured at three time points (3-, 6- and 12-months post-injury) across one year. To do so, we used a publicly available resting state fMRI dataset of patients with moderate-to-severe TBI (N=14 in each session) and age-matched healthy individuals (N=12) and extracted the time series from each of the 1,000 parcels in the fine-grained Schaefer parcellation (Schaefer et al., 2018). We tested the hypotheses that whole-brain amplitude turbulence would (i) accurately differentiate patients with TBI from healthy controls (HCs) and (ii) progressively approach the values of HCs across the one-year recovery trajectory. Furthermore, we expected to see other measures derived from the turbulent dynamics framework to discriminate between TBI patients and HCs. Finally, we anticipated that the whole-brain computational model would demonstrate that the brain’s capability to respond to external *in silico* perturbations is compromised after TBI.

a. 2. **Methods**

### 2.1. Model-free approach

#### 2.1.1. Participants and behavioral data

The source of data for this study (Fig.1a) was obtained from the open access repository OpenNeuro (https://openneuro.org/datasets/ds000220/versions/00002). Please refer to the original publication (Roy et al., 2017) for a full description of the participants included. Briefly, 14 patients with moderate-to-severe TBI between the ages of 18 and 36 with education ranging from 12 to 18 years, and 12 healthy controls (HCs) of comparable age and education were included. A Glasgow Coma Scale (GCS) at time of injury was indicative of severe (3-8) and moderate (9-12) injury. The NIfTI images from 2 TBI patients were discarded due to incorrect labels (superior-inferior swap) as shown in FSLeyes. All subjects with TBI completed three separate sessions (3-, 6-, and 12-months following injury). Resting state fMRI (rs-fMRI) data from HCs at baseline was also included for group comparisons. The authors of the original publication also provided the *z* scores for the neuropsychological battery used to evaluate TBI patients at 3-months post-injury. These included the Visual Search and Attention Test (VSAT), WAIS-III Digit Span (Forward, Digits-F and Backward, Digits-B), Trail Making Test B (TMT), and Stroop Color and Color-Word (Stroop).

#### 2.1.2. Resting state fMRI acquisition

Five-minute rs-fMRI data were acquired using a whole-brain EPI sequence (150 volumes, TR=2000 ms, TE=30 ms, flip angle=90°, 240 × 240 mm field of view, 80 × 80 acquisition matrix, voxel size 1.9 x 1.9 x 4 mm). In addition, a high-resolution whole-brain T1-weighted image was acquired (voxel size 1mm^3^).

#### 2.1.3. Resting state fMRI preprocessing

Preprocessing was conducted using Statistical Parametric Mapping software (SPM12, Wellcome Department of Cognitive Neurology, University College London) running under Matlab Release 2021a (The MathWorks, Inc., Natick, MA). The first 5 volumes were removed to control for signal instability. Preprocessing steps included slice timing correction, realignment, segmentation, functional to structural coregistration, spatial normalization (voxel size 2mm^3^), and smoothing (8mm Gaussian kernel).

We then applied a denoising pipeline in order to minimize the variability due to physiological, outlier, and residual subject-motion effects (Nieto-Castanon, 2020). The preprocessed output files from SPM were imported into the CONN toolbox version 20.b (https://www.nitrc.org/projects/conn), including the following subject- and session-specific files: T1 and rs-fMRI scans; segmented grey matter (GM), white matter (WM), and cerebrospinal fluid (CSF) images; and 6 realignment parameters after rigid body motion correction. We applied the same pipeline as in our previous publication with TBI patients (Martínez-Molina et al., 2022; see Supporting Information for more details). The 1,000 nodes and 400 nodes Schaefer parcellation in MNI152 volumetric space were added as an atlas in the CONN toolbox to extract the timeseries from each node (Schaefer et al., 2018). The Euclidean distance between parcels’ centroids from the Schaefer parcellation was also calculated as in our previous publications (Deco & Kringelbach, 2020; Escrichs et al., 2022).

#### 2.1.4. Probabilistic tractography analysis

We used the diffusion-weighted and T2-weighted neuroimaging data from 32 participants of the HCP database, as previously reported (Deco & Kringelbach, 2020). In the HCP website, a description of the acquisition parameters is detailed (Setsompop et al., 2013). We used the Lead-DBS software package (https://www.lead-dbs.org/) for the preprocessing of tractography data (Horn et al., 2017) which, briefly, were processed by using a q-sampling imaging algorithm implemented in DSI studio (http://dsi-studio.labsolver.org). Segmentation of the T2-weighted anatomical images and co-registering the images to the b0 image of the diffusion data using SPM12 produced a WM mask. For each individual, 200,000 fibers were sampled within the WM mask. Fibers were transformed into MNI space using the Lead-DBS software (Horn & Blankenburg, 2016). Finally, the structural connectivity (SC) using the Schaefer 1,000 parcellation (Schaefer et al., 2018) was extracted with the standardized methods in the Lead-DBS. The SC was averaged across participants before the simulated attack.

#### 2.1.5 Simulated attack

In this approach, we used the binary lesion maps from the TBI patients provided by the authors of the original publication (Roy et al., 2017) in order to simulate connectome attacks relative to the SC derived from healthy participants (see previous section). Our goal was to create a lesion mask matrix based on the lesions in the TBI patients that could be applied to the healthy SC. A similar methodology has been recently implemented in patients with aphasia and provided a good approximation of the effects of the real lesions on anatomic networks as well as explaining a comparable amount of variance in behavioral data (Medaglia et al., 2022). Further, virtual lesions have allowed to disentangle the local and recurrent components from TMS-EEG evoked potentials using a whole-brain connectome-based computational modeling (Momi et al., 2023).

The lesion masks for each TBI patient (N=11, one patient did not have visible lesions) in native space were normalized to MNI space (after normalization one patient was discarded due to misalignment) and the overlap with each node from the Schaefer 1000 nodes parcellation computed using the fslstats command in FSL v.6.0 (https://fsl.fmrib.ox.ac.uk/fsl/). This value was divided by the volume of the Schaefer node to account for variability in the size of the nodes. We created the lesion mask matrix to simulate the lesion on the healthy SC using 4 thresholds: 1.5, 2, 3 and 4 standard deviations (SD) from the mean overlap volume of lesions in all TBI patients. Thus, for each TBI patient, we calculated the nodes to attack based on these four thresholds. This allowed us to calculate the frequency of lesion for a given node as well as find the unique nodes to attack, that is, nodes that were lesioned in any given patient. The lesion mask matrices thresholded at 3 and 4 SD only resulted in 10 and 7 nodes to attack and were discarded. The lesion mask matrices thresholded at 1.5 and 2 SD produced 41 and 22 nodes to attack. The simulated attack with the 1.5 and 2 SD lesion matrices was performed in two ways leading to the disconnection of the lesioned node: (1) a complete deletion of the SC between the node and the rest of the brain (binary approach, value set to 0 in the lesion mask matrix); (2) a weighted downregulation of the SC between the node and the rest of the brain. The weight was computed by calculating the number of patients with the lesioned node divided by the total number of TBI patients. Then the value in the lesion mask matrix between that node and the rest of the brain was set to 1 - weight. For each masking threshold, the SC was multiplied by the lesion mask matrix before running the computational modeling. We took a dual approach, one binary (1) and one weighted (2), in order to ascertain the sensitivity of the whole-brain model perturbational measures to the effect of virtual disconnections. A list of the nodes to attack for the 1.5 SD threshold, the corresponding anatomical region and the frequency of lesion can be found in Table S19.

#### 2.1.5. Amplitude turbulence

We measured amplitude turbulence by first defining the Kuramoto local order parameter and then taking the standard deviation of the modulus across time and space. First, we defined the amplitude turbulence, R_λ_ (𝑥̅,t), as the modulus of the Kuramoto local order parameter for a given brain region as a function of the time at each spatial scale λ:

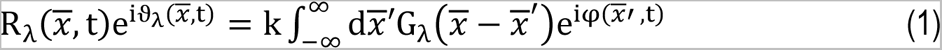

where φ(𝑥, t) denotes the phases of the BOLD signal data, G_λ_ refers to the local weighting kernel G_λ_(𝑥̅) = e^−λ|𝑥̅|^ and k denotes the normalization factor 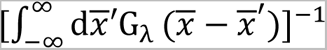. The BOLD time series were filtered with a second-order butterworth filter in the range between 0.008-0.08 Hz, where we chose the typical high pass cutoff to filter low-frequency signal drifts (Fox et al., 2005; Glerean et al., 2012) and the low pass cutoff to filter the physiological noise, which tends to dominate the higher frequencies (Fox et al., 2005; Glerean et al., 2012). We then applied the Hilbert transform in order to obtain the complex phase of the BOLD signal for each brain node as a function of time.

Local levels of synchronization at a certain λ, as a function of space (𝑥̅) and time (t) are defined by R_λ._ We then measure the amplitude turbulence, D, as the standard deviation across nodes and time of the modulus of the Kuramoto local order parameter (R_λ_):

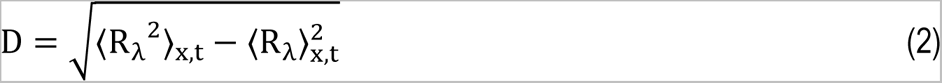

where the brackets 〈〉_x,t_ denote averages across space and time.

This measure captures the brain vortex space over time, motivated from fluid dynamics and the rotational vortices (Deco & Kringelbach, 2020). Specifically, we explored the level of Kuramoto amplitude turbulence over different λ values, i.e., from 0.01 (∼100 mm) to 0.30 (∼3 mm), in steps of 0.03.

#### 2.1.6. Information cascade flow

The information cascade flow indicates how the information travels from one scale (λ) to a lower scale (λ − Δλ, where Δλ corresponds to a scale step) in successive time steps (t and t + Δt). It is calculated as the temporal correlation between the Kuramoto local order parameter in two successive scales and times as:

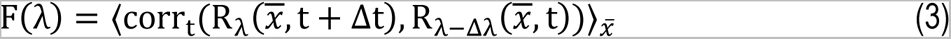

where the brackets 〈〉_𝑥̅_ indicate averages across time and nodes. Then the information cascade is obtained by averaging the information cascade flow across scales λ, which captures the whole behavior of the information processing across scales.

#### 2.1.7. Information transfer

The information transfer captures how information travels across space at each λ scale. It is calculated as the slope of a linear fitting, in the log-log scale, of the temporal correlation between the Kuramoto local order parameter of two nodes at each scale as a function of its Euclidean distance (r) in the inertial subrange.

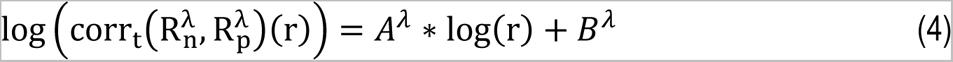

where 𝐴^𝜆^ and 𝐵^𝜆^ are the fitting parameters for the λ scale, and r denotes the distance in the brain.

#### 2.1.8. Local node-level metastability

The node-level metastability, NLM, defines the variability of the local synchronization of the nodes and is computed as the standard deviation across time of the Kuramoto local order parameter as follows:

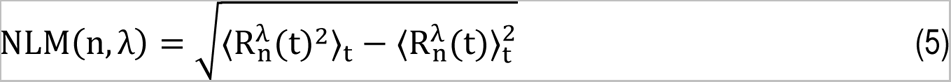

where brackets 〈〉_t_ denote average across time.

Here, we estimated the discrete version of the node-level Kuramoto order parameter, with modulus R and phase ν, which represents a spatial average of the complex phase factor of the local oscillators weighted by the coupling as:

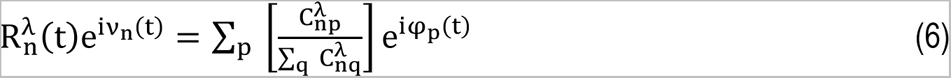

where ϕ_p_(t) are the phases of the spatiotemporal data and 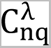 is the local weighting kernel between node n and q, and λ defines the spatial scaling:

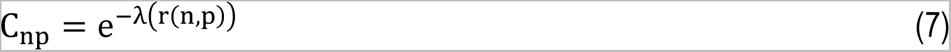

### 2.2. Model-based approach

#### 2.2.1. Whole-brain Hopf model

We built whole-brain dynamical models with Stuart-Landau oscillators based on the normal form of a supercritical Hopf bifurcation (Deco et al., 2017a). This type of bifurcation can change the qualitative nature of the solutions from a limit cycle that yields self-sustained oscillations toward a stable fixed point in phase space. This model is characterized by model parameters that rule the global dynamical behavior. One control parameter is the multiplicative factor (G) representing the global conductivity of the fibers scaling the structural connectivity between brain areas, which is assumed to be equal across the brain (Deco et al., 2017a,b). The other relevant parameters are the local bifurcation parameter (an), which rules the dynamical behavior of each area between noise induced (a < 0), self-sustained oscillations (a > 0), or a critical behavior between both (a ∼ 0) (Fig.1b). The model parameters were optimized to better fit the empirical functional connectivity as a function of the distance, r, within the inertial subrange. The models consisted of 1,000 brain regions from the resting state atlas (Schaefer et al., 2018). The underlying structural connectivity matrix C_np_ was added to link the brain structure and functional dynamics by computing the exponential distance rule (Equation 7). The local dynamics of each brain region were characterized by the normal form of a supercritical Hopf bifurcation, which emulates the dynamics for each region from noisy to oscillatory dynamics as:

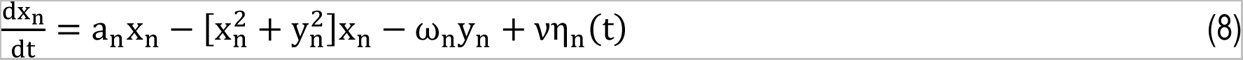

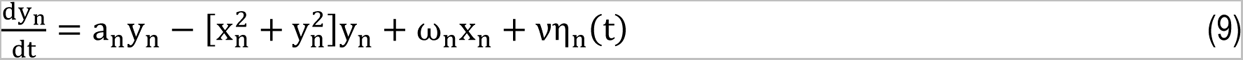

where η_n_(t) is the additive Gaussian noise with standard deviation ν = 0.01. This normal form has a supercritical bifurcation at a_n_ = 0, such that for a_n_ > 0, the system is in a stable limit cycle oscillation with frequency f_n_ = ω_n_/2π, while for a_n_ < 0 the local dynamics are in a stable point (the noisy state). The frequency ω_n_ of each region was obtained from the empirical functional MRI data as the power spectrum peak.

Finally, the whole-brain dynamics were determined by the set of coupled equations as follows:

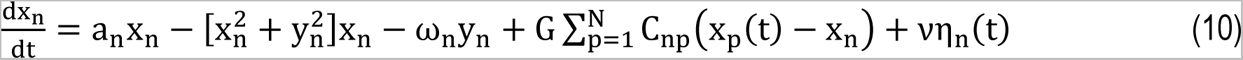

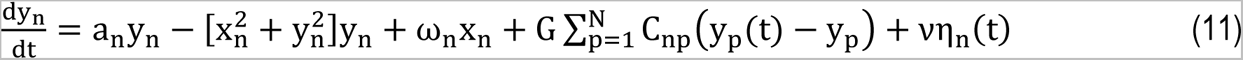

where 𝐶_np_ is the structural connectivity weight between node 𝑛 and 𝑝, and 𝐺 is the global coupling weight with equal contribution between all the nodal pairs. The C_np_ was derived using the exponential distance rule and long-range exceptions (Deco et al., 2021). We used λ=0.18 as this was shown to produce the best fit. This parameter is different from the λ values used to calculate global amplitude turbulence at different spatial scales. For each simulated attack (4 in total: 1.5 and 2 SD, binary and weighted approach), this C_np_ was multiplied by the corresponding lesion mask matrix. The output of the model is a set of complex-valued BOLD-like signals with time-varying intrinsic frequencies (ω). The real part is considered to represent BOLD signal recorded in the fMRI session whereas the imaginary part characterizes the hidden state of the oscillator unseen to the scanner.

#### 2.2.2. Functional connectivity fitting

The functional connectivity fitting of the BOLD signal data was assessed by applying Kolmogorov’s structure-function. The structure-function for a variable u characterizes the evolution of the functional connectivity (FC) as a function of the Euclidean distance between equally distant brain regions and is defined as follows:

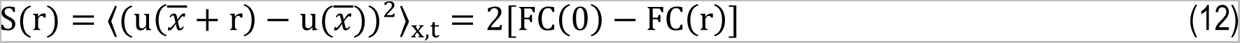

where FC(r) is the spatial correlations of two points separated by a Euclidean distance r, and is given by:

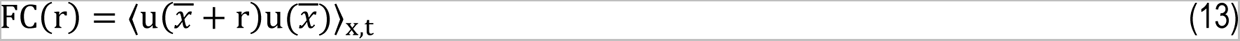

where 〈〉_x,t_ denotes average across the spatial x coordinates of the nodes and time. Then, we calculated the fitting as the Euclidean distance between simulated and empirical FC(r) within the inertial subrange (Deco & Kringelbach, 2020).

#### 2.2.3. Susceptibility and Information encoding capability

The susceptibility measure obtains the brain’s sensitivity to react to external stimulation. The Hopf model was perturbed for each G by randomly changing the local bifurcation parameter, a_n_, in the range [-0.02:0] and was computed by measuring the modulus of the Kuramoto local order parameter as:

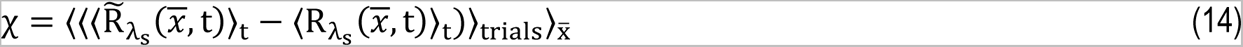

Where 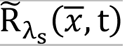 corresponds to the perturbed case, the R_λs_(𝑥̅, t) to the unperturbed case, and 〈〉_t_, 〈〉_trials_ and 〈〉_𝑥̅_ to the average across time, trials, and space, respectively.

The Information encoding capability (I) captures how the external stimuli are encoded in whole-brain dynamics. This measure was defined as the standard deviation across trials of the difference between the perturbed 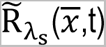 and unperturbed R_λs_(𝑥̅,t) mean of the modulus of the local Kuramoto order parameter across time t, averaged across all brain regions n as follows:

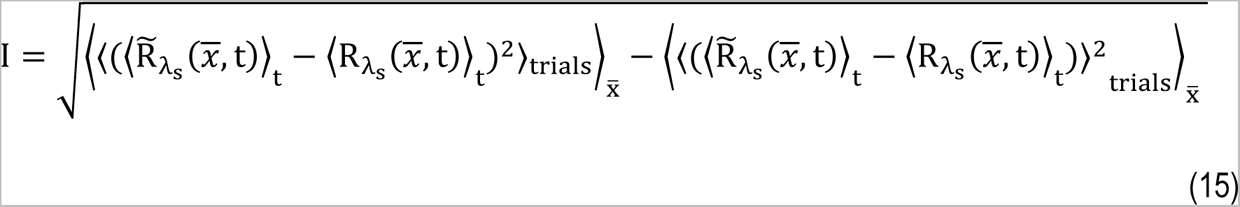

where the brackets 〈〉_t_, 〈〉_trials_ and 〈〉_x̅_ denote the averages defined as above.

### 2.3. Statistical analyses

A 4 (Group) x 10 (Lambda) ANOVA was used to compare TBI patients and controls for global amplitude turbulence and information transfer. Information cascade flow differences were assessed with a 4 (Group) x 9 (Lambda) ANOVA. One-way ANOVA (4 groups) was used to test significant differences in information cascade. In all cases, post hoc tests were performed to evaluate pairwise differences. The longitudinal effects within the TBI group were assessed with repeated measures ANCOVA with lesion volume as nuisance covariate followed by post hoc comparisons. Differences in RSN turbulence were assessed with one-way ANOVA (4 groups) followed by post hoc tests. Differences in node-level metastability were examined with the Kolmogorov-Smirnov test. Pearson’s correlation were performed to assess brain-behavior correlations. A Wilcoxon signed rank test was used to determine differences within HCs or TBI groups in susceptibility and information encoding capability. Between groups differences were assessed with a Wilcoxon ranked sum test. When needed, we corrected for multiple comparisons using false discovery rate (FDR) with the method of Benjamini and Hochberg (Benjamini and Hochberg, 1995).

## 3. Results

Our findings are split into two different approaches: model-free and model-based. In the first approach, we computed empirical measures of whole-brain turbulent dynamics using a fine-grained parcellation with 1,000 regional phase time courses derived from resting-state fMRI in both HCs (N=12) and patients with TBI (N=12) at 3-, 6- and 12-months post-injury (see Fig.1a and Material and Methods for more details). Since there were not significant differences between session 1 and 2 in HCs (see Table S1), we averaged the values for both sessions. In the model-based approach, we built a whole-brain dynamical model for each condition and group to explore the sensitivity of TBI patients and HCs to react to external *in silico* perturbations. For each whole-brain model, we applied *in silico* perturbations by changing the local bifurcation parameter and computed the susceptibility and information encoding capability measures. A whole-brain model was computed separately for each stage of recovery in TBI patients and for each simulated attack approaches as well as with the intact structural connectivity (see Fig.1b and Material and Methods for more details). Given the significant differences between session 1 and 2 in HCs for the perturbational measures (Table S1), these were not averaged across sessions.

### 3.1 Model-free measures of synchronization

#### 3.1.1 Empirical measures of turbulence are reduced after traumatic brain injury

TBI patients showed differences compared to HCs for each turbulent measure. To examine global amplitude turbulence differences between HCs and TBI patients at the 10 spatial scales under study, we used a 4 (Group) x 10 (Lambda) ANOVA. The results from this analysis revealed a significant Group x Lambda interaction (F_(27,440)=_2.534, p<0.0001, Table S2, Fig.2a). Post hoc tests indicated that HCs’ amplitude turbulence was significantly higher than that of TBI patients across all stages of recovery for λ=0.01 and λ=0.03 (Table S2), which are the scales including larger spatial distances across nodes. For λ=0.06, we also found significant differences between HCs and TBI patients at 3- and 6-months post-injury (Table S2). It is worth noting that the greatest mean difference was observed between HCs and TBI patients at 6-months at all significant λ values. The rest of λ values did not show any significant difference between HCs and TBI patients post hoc.

The information cascade flow also captured differences in information propagation between consecutive spatial scales. We found a significant Group x Lambda interaction (F_(24,396)=_3.007, p<0.0001, Fig.2b). Like the findings with amplitude turbulence, post hoc tests revealed group differences at larger spatial scales. Specifically, all group comparisons were significant for λ=0.01 and between HCs and TBI at 6-months for λ=0.03 (Table S3). In the one-way ANOVA, we found a significant group effect for information cascade (F_(3,44)=_7.061, p<0.001, Fig.2c). As indicated by post hoc tests, a significant group difference was found between HCs and TBI at 3- (T_(22)_=2.833, p=0.034, Table S4) and 6-months (T_(22)_=4.552, p<0.001, Table S4).

The Group x Lambda interaction was also significant for information transfer (F_(27,440)=_2.325, p<0.001, Fig.2d). In this case, however, differences were found at larger and intermediate λ values (λ=0.27-0.15), with all group comparisons differing for λ=0.27 and λ=0.24 (Table S5).

We repeated these analyses with the 400 nodes Schaefer parcellation. Although the pattern of results regarding the spatial scales affected was not identical, the results also indicated that turbulent measures were systematically decreased in TBI patients as compared to HCs (see Tables S6-9).

In an additional analysis, we assessed longitudinal differences within TBI patients and the potential confounding effect of lesion volume by using a repeated measures ANCOVA with time points as within-group factor and lesion volume as nuisance covariate. We found a significant Group x Lambda interaction in global amplitude turbulence (F_(18,180)_=2.073, p=0.008, Fig. S1a). Post hoc comparisons revealed that global amplitude turbulence was significantly decreased after 6-months compared to 3-months post-injury for λ=0.01 (T(22)= 3.844, p=0.018, Table S10). Information cascade flow also showed a significant Group x Lambda interaction (F_(16,160)_=1.809, p=0.034, Fig. S1b). Post hoc, we found a significant reduction at 6-months post-injury stage compared to 3-months for λ=0.01 (T_(22)_=4.733, p<0.001, Table S11). Information cascade was significantly reduced at the 6-months post-injury stage compared to 12-months (group effect: F_(2,20)_ = 4.205, p = 0.003; T_(22)_=2.906, p=0.026, Fig.S1c,Table S12). No significant differences were found for information transfer in the within-group analysis (Table S13).

#### 3.1.2 Reductions in RSN empirical turbulence at the 6-month stage show recovery at the 12-month stage

Next, we examined differences in amplitude turbulence between HCs and TBI patients in the 7 RSNs included in the Schaefer parcellation for the spatial scales where we found significant reductions in global amplitude turbulence between HCs and TBI patients at all stages of recovery (λ=0.01 and λ=0.03, see previous section). The results of the one-way ANOVA including amplitude turbulence in HCs (averaged across both sessions) and TBI patients at 3-, 6- and 12-months post-injury revealed a significant group effect for λ=0.03 in all 7 RSNs except for the limbic network (Table S14, Fig.3a). Post hoc we found that this effect was in all cases attributable to the difference between HCs and TBI patients at 6-months post-injury. In other words, this implies that after a decreased in turbulent dynamics at 6-months most RSNs recovered healthy levels by 12-months at this spatial scale. No significant group effect was found for λ=0.01.

To complement this analysis, we computed the Kolmogorov-Smirnov distance (KSD) that quantifies the difference in the distributions of node-level metastability between HCs and TBI patients. We found that the KSD monotonically decreases (i.e., distributions are more similar) across scales in all group comparisons, whereas the value of λ increases. In other words, the KSD is maximal for lower values of λ, i.e., long distances in the brain (Fig. 3b). Remarkably, the highest KSD is found between HCs and TBI patients at 6-months post-injury (Table S15).

**Figure 1:**
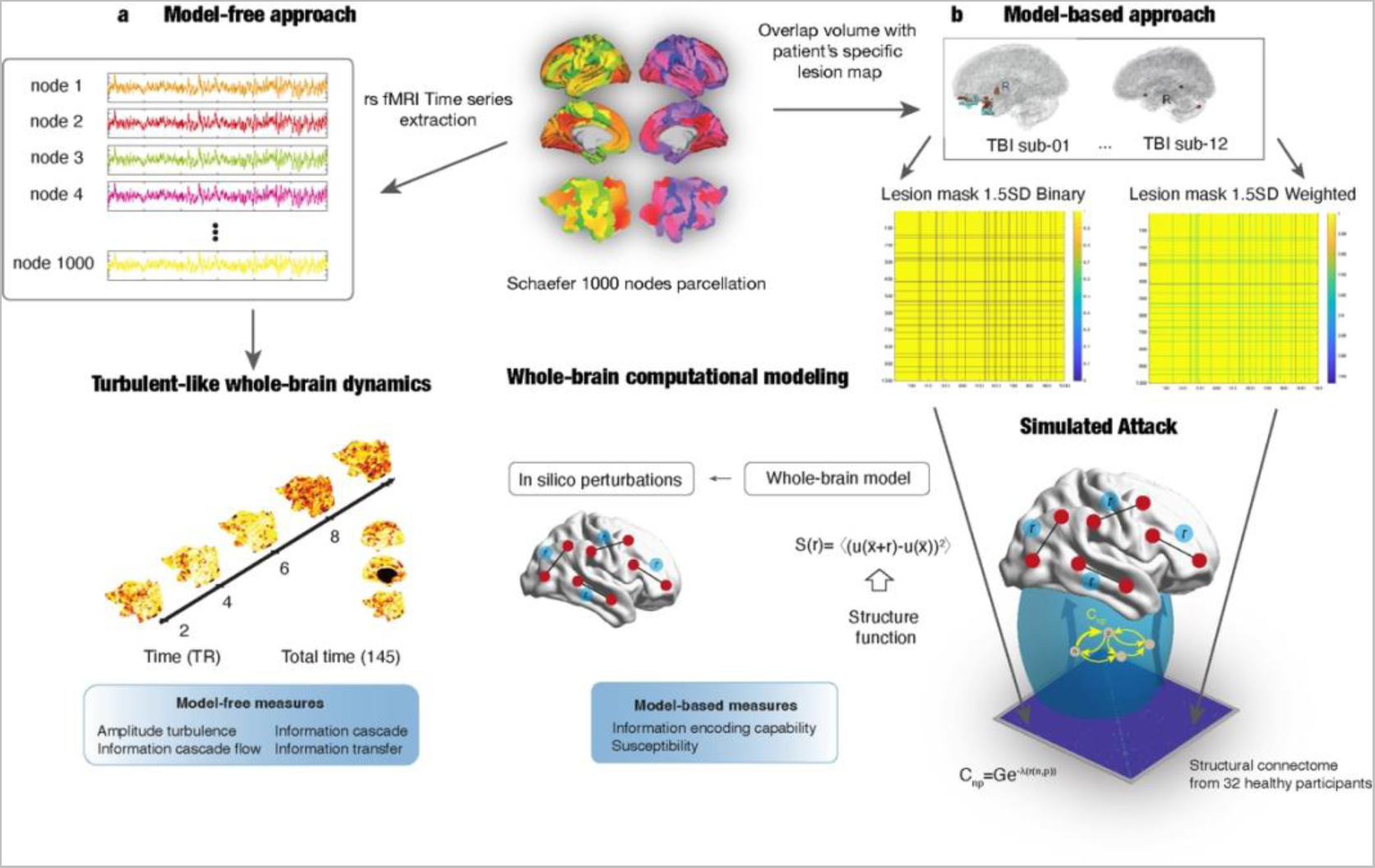
Schematic of the model-free and model-based analysis pipelines. We used an open access longitudinal dataset from a previous publication (Roy et al., 2017). T1-weighted (T1-W) and resting state fMRI (rs-fMRI) data from traumatic brain injury (TBI) patients were collected at three time points (3-, 6- and 12-months post-injury). For group comparisons, T1-W and rs-fMRI were collected from age-matched healthy controls in two sessions. **(a)** Model-free approach. Top left: rs-fMRI data were preprocessed in the CONN toolbox and the time series were extracted for each of the 1,000 nodes in the Schaefer parcellation and the instantaneous phase calculated (Schaefer et al, 2018). Bottom left: Visualization of the change over time and space of the Kuramoto local order parameter in a healthy subject (left hemisphere). The Kuramoto local order parameter can define the turbulent signatures of brain activity at different spatial scales in the vortex space. Here, we computed four turbulent measures based on the Kuramoto local order parameter to characterize the brain’s information processing: amplitude turbulence, information cascade flow, information cascade and information transfer (see Methods for more details). **(b)** Model-based approach and simulated attack. We built Hopf whole-brain dynamical models that described the intrinsic dynamics of each brain area by a Stuart-Landau non-linear oscillator. Such local oscillators are linked through the underlying structural connectivity to simulate the global dynamics. We used the DTI matrix with the exponential distance rule and long-range exceptions as structural connectivity to fit the empirical data as a function of the Euclidean distance. To account for the effect of focal lesions, we adapted a simulated attack approach (Medaglia et al., 2022). The overlap between the patient’s specific lesion mask and the Schaefer parcellation was calculated and used to create a group lesion mask thresholded at 1.5 and 2 SD from the group’s overlap. Next, a binary or weighted disconnection was applied to the group lesion mask. For each threshold and disconnection approaches, a model was computed by multiplying the structural connectivity by the group lesion mask. Then we applied *in silico* perturbations to assess the model’s reaction for each time point in the TBI group compared to healthy controls using the susceptibility and information encoding capability model-based measures (bottom panels adapted with permission from Deco & Kringelbach, 2020).

**Figure 2:**
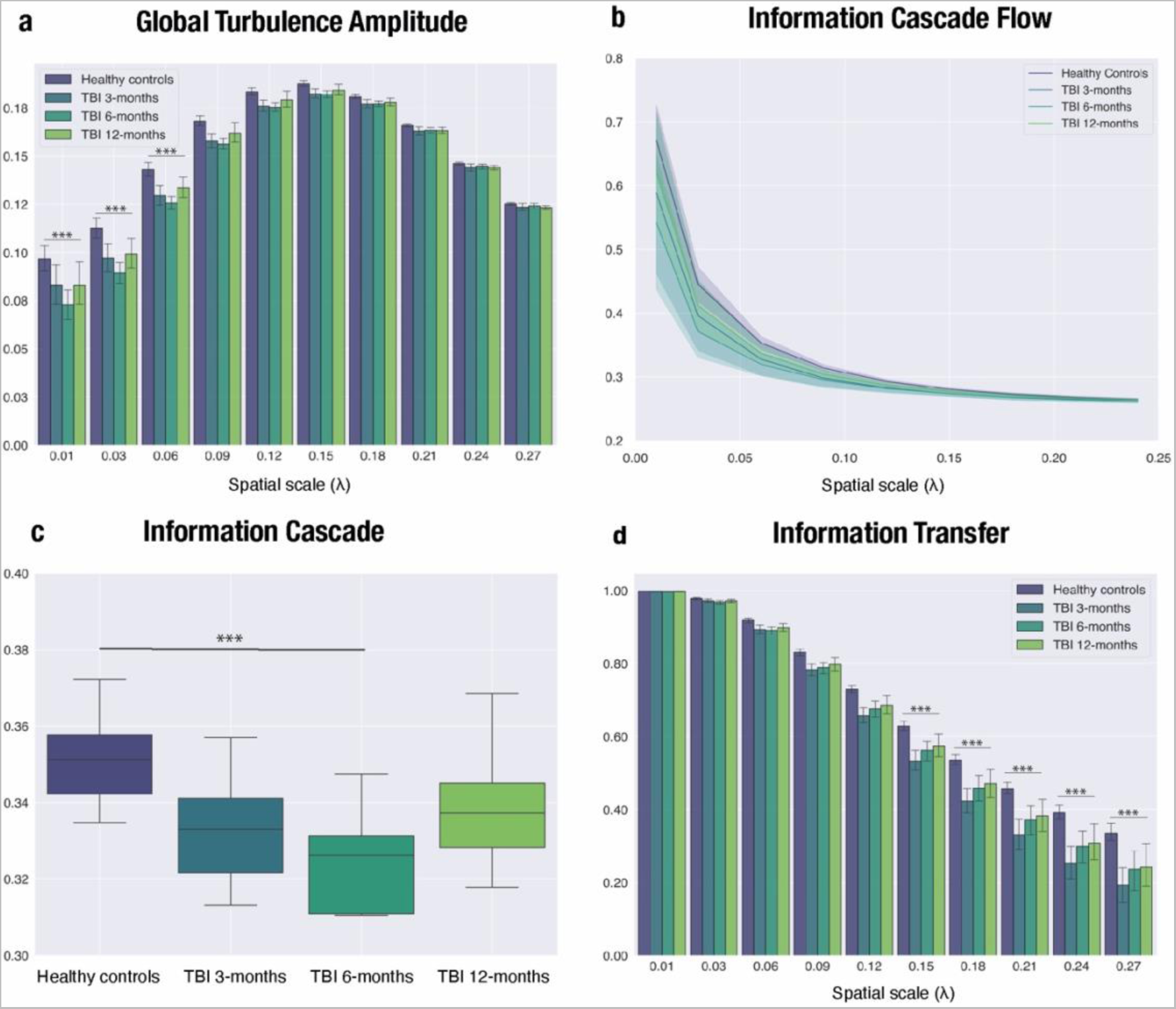
Turbulent dynamic measures differ between TBI patients and healthy controls over time, peaking at 6-months post-injury. Significant differences in turbulent-like dynamics between healthy controls (HCs, averaged across sessions) and TBI patients at 3-, 6- and 12-months post-injury were calculated using a 4 (Group) x 10 (Lambda) ANOVA. The Group x Lambda interaction was significant at long distances both for global amplitude turbulence (λ=0.01-0.06) **(a)** and information cascade flow (λ=0.01-0.03) **(b).** We found a group effect for information cascade **(c).** Post hoc tests revealed that it was due to the difference between HCs and TBI patients at 6-months post-injury. **(d)** For information transfer, the Group x Lambda interaction was significant for intermediate and short distances (λ=0.01-0.03). Bar plots depict mean values (+SEM). The lower and upper edges of the boxplots indicate the first (25th percentile) and third quartile (75th percentile), middle lines are medians. Asterisks represent the effect of the interaction for all measures except for information transfer where the results from the post hoc test are shown (***p <0.001-0.001).

**Figure 3:**
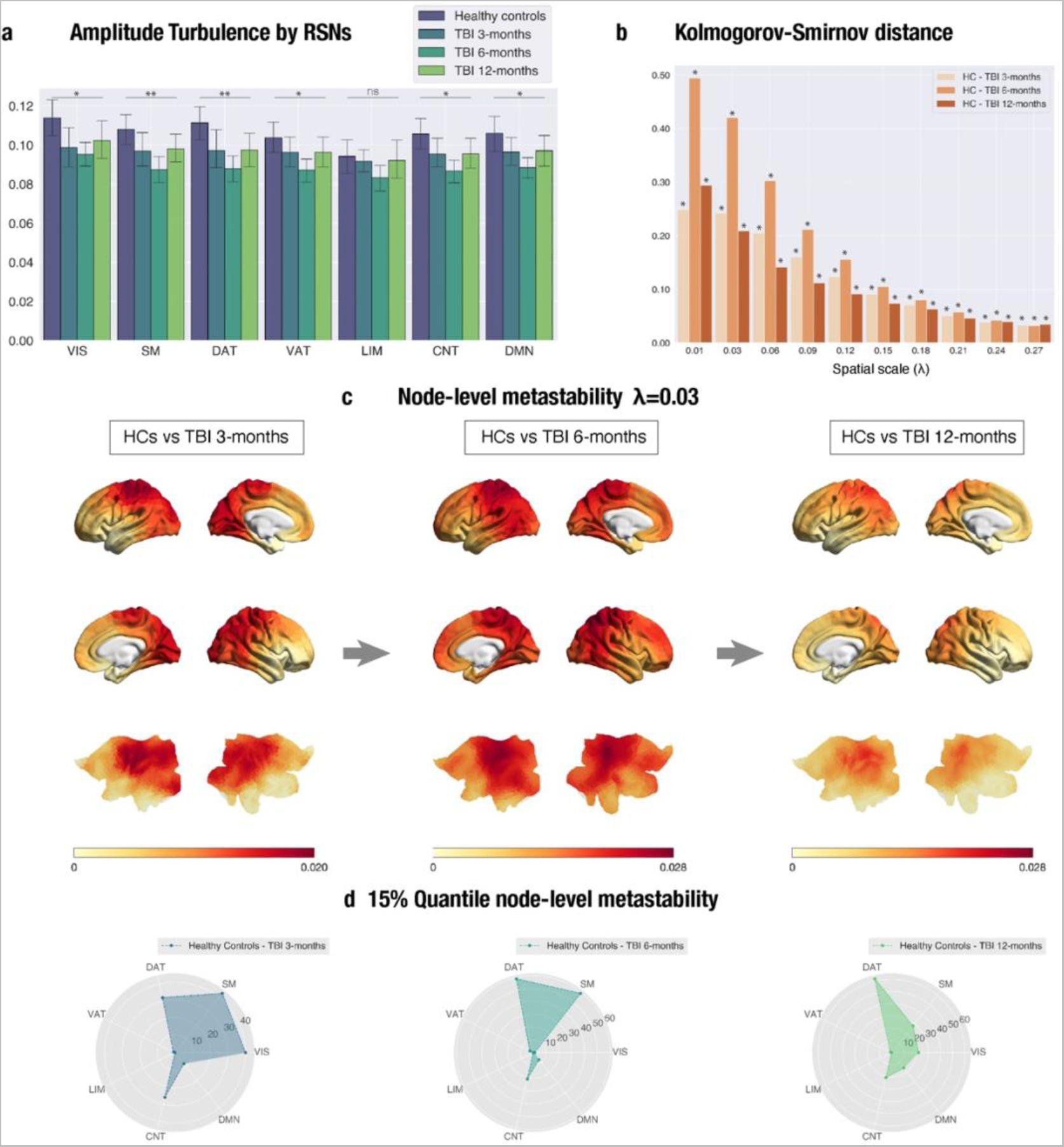
Amplitude turbulence in most RSNs shows a U-shape recovery trajectory with no significant differences between HCs and TBI patients by 12-months post-injury (λ=0.03). Nodes in the somatosensory, dorsal attention and control networks show the highest difference in TBI patients at 6-months post-injury. (a) Amplitude turbulence is shown for each dataset and the 7 RSNs included in the Schaefer 1,000 nodes parcellation. Asterisks indicated significance level for the group effect in the comparison with healthy controls (HCs, averaged across sessions). (b) Kolmogorov-Smirnov distance (KSD) distributions for the difference in node-level turbulence between HCs (averaged across sessions) and TBI patients at 3- (left), 6- (middle) and 12-months post-injury. Of note, the largest KSD values were reached for lower λ values (large distances) at the 6-months post-injury stage for all spatial scales (λ). (b) Render brains represent the absolute difference of the node-level metastability between each dataset for scale λ = 0.03. (d) Radar plots representing the number of nodes on the top 15% quantile of the absolute difference for the HCs vs TBI patients at 3-, 6- and 12-months post-injury comparison for λ =0.03. At 6-months post-injury, most of the nodes were ascribed to the SM, DAT and CNT networks. Abbreviations: LH: left hemisphere; LIM, limbic; CNT, control; DMN, default mode; DAT, dorsal-attention; VAT, ventral attention; VIS, visual; SM, somatosensory. Asterisks: *p <0.05, **p <0.01 (RSN turbulence); * denotes significant differences (KSD, for specific p-values see Table S14).

**Figure 4.**
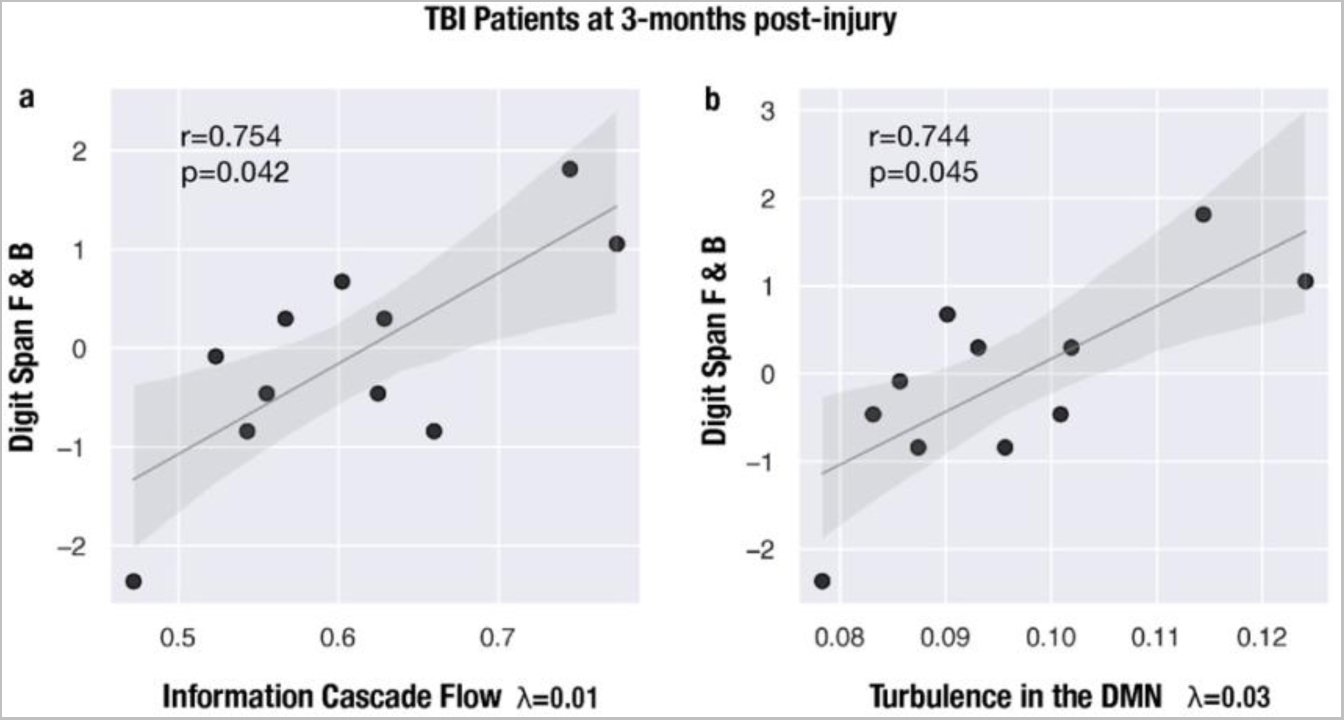
Working memory performance is positively associated with turbulent measures in TBI patients at 3-months post-injury. Scatter plot representing the positive correlation between *z* scores in the WAIS-III Digit Span (Forward and Backward) and information cascade flow for λ=0.01 **(a)** and turbulence in the DMN for λ=0.03 (**b).** Abbreviations: F: forward, B: backward, DMN: Default Mode Network.

For illustrative purposes, Fig. 3c displays the absolute difference in node-level metastability between HCs and TBI patients at 3-,6- and 12-months post-injury at λ= 0.03 rendered onto the brain cortex. Additionally, we computed the number of nodes on the top 15% quantile of the absolute difference in node-level metastability for the HCs vs TBI patients at 6-months post-injury comparison (Fig. 3d). The nodes with the largest differences in node-level metastability were located in the dorsal attention, somatomotor and control networks.

#### 3.1.3 Empirical measures of turbulent dynamics predict cognitive performance

To determine whether empirical measures of turbulent dynamics relate to cognitive impairment in TBI patients, we performed Pearson’s correlations with the neuropsychological battery of tests and turbulent measures at 3-months post-injury as provided by the authors of the original publication. The battery included those tests more sensitive to detect deficits in processing speed and working memory that are common after TBI (see Material and Methods). We found a positive association between Digital Span (Forward and Backward) and information cascade flow at λ=0.01 (large spatial scale) (r=0.754, p=0.007, Fig.4a, Table S16). This result survived multiple-comparison correction with FDR across number of battery tests (p-adj=0.042). In other words, the greater the propagation of information between consecutive spatial scales, the better the working memory performance. Furthermore, we found that performance in this working memory task was also positively associated with turbulence in the default mode network for λ=0.03 (r=0.744, p=0.009, Fig.4b, Table S17). This result also survived multiple-comparison correction with FDR across number of battery tests (p-adj=0.045). Lesion volume did not correlate neither with cognitive performance nor with amplitude turbulence.

### 3.2 Model-based framework

We defined a whole-brain model of coupled oscillators (Stuart-Landau oscillators) for each time point of the TBI patients and HCs rs-fMRI data. To account for the effect of focal lesions on the structural connectivity (SC) we adopted a simulated attack approach with lesion mask matrices thresholded at 1.5 and 2 standard deviations (SD) and applied a binary or weighted attack resulting in a total of 4 simulated attacks (see Fig.1b). Additionally, for each time point of the TBI patients’ dataset, a computational model was run with intact healthy SC to assess the sensitivity of the model to the simulated attack. In total, 11 different whole-brain computational models were computed for the 1.5 SD lesion mask. To optimize the fitting between the whole-brain model and the empirical rs-fMRI data, we varied the global coupling parameter G from 0 to 3 in steps of 0.01, and for each G value, we ran 100 simulations with the same TR (2s) and time duration (145 scans) as the empirical rs-fMRI data. We then determined the optimal working point of each model as the minimum of the fitting level. Next, we calculated differences in susceptibility and information encoding capability and susceptibility between groups (HCs session 1 and 2; TBI patients at 3-, 6- and 12-months post-injury) after *in silico* perturbations.

We first identified the working point G for each rs-fMRI dataset, which represents the global coupling parameter. First, we found that the working point of the model was lower for the TBI patients at all time points (3-,6- and 12-months post-injury) as compared with the value for both sessions of HCs (Fig.5a-c) regardless of whether an intact or lesioned SC was used as input to the model. Crucially, we detected the same pattern, that is, the global coupling parameter for TBI patients at 6-months was smaller than that at 3- and 12-months post-injury. Second, the global coupling parameter shifted to lower values after the simulated attack with both thresholds (1.5 and 2 SD). However, the difference with respect to the intact healthy SC was small. And the binary approach did not yield to smaller values than the weighted (see Table S18).

Once we identified the working point G for each rs-fMRI dataset, we introduced external perturbations by changing the local bifurcation parameter a_n_ of each brain node n. For each value of G, we perturbed the whole-brain model 100 times with random parameters for the local bifurcation parameter. Our model-based measures, susceptibility and information encoding capability, allowed us to evaluate the reactivity to the external perturbations introduced (for more details see Material and Methods). Briefly, susceptibility was estimated by first measuring the perturbed and non-perturbed modulus of the Kuramoto local order parameter and then calculating the difference between the perturbed and non-perturbed cases averaged across time, trials, and brain nodes. As an extension of this measure, the information encoding capability of the whole-brain model was calculated as the standard deviation across trials of the difference between the perturbed and unperturbed mean of the modulus of the local order parameter across time, averaged over all brain nodes.

Fig. 5d-e shows that both susceptibility and information encoding capability were significantly reduced in TBI patients across the recovery trajectory compared to both sessions of HCs regardless of whether an intact or lesioned SC was used as input to the model. However, the values of these measures for the binary simulated attack were significantly reduced at all stages of recovery with respect to the values in the weighted simulated attack suggesting that the model is most sensitive to focal lesions that greatly overlap. Since in this cohort of TBI patients the frequency of lesion per node was mostly 1 (Table S19), the results from the weighted attack may be more representative of the underlying average SC. With the 1.5 SD weighted lesion mask matrix, TBI patients at 6-months post-injury showed the lowest mean value across trials (Fig.5d-e, Tables S20-21). This pattern of results was replicated when using the weighted 2 SD lesion mask (Fig. S2, Tables S22-23). Therefore, the results from our *in silico* stimulation suggest that, at 6-months post-injury, the brain in this cohort of TBI patients is working in a regime that is the least responsive to external perturbations.

**Figure 5:**
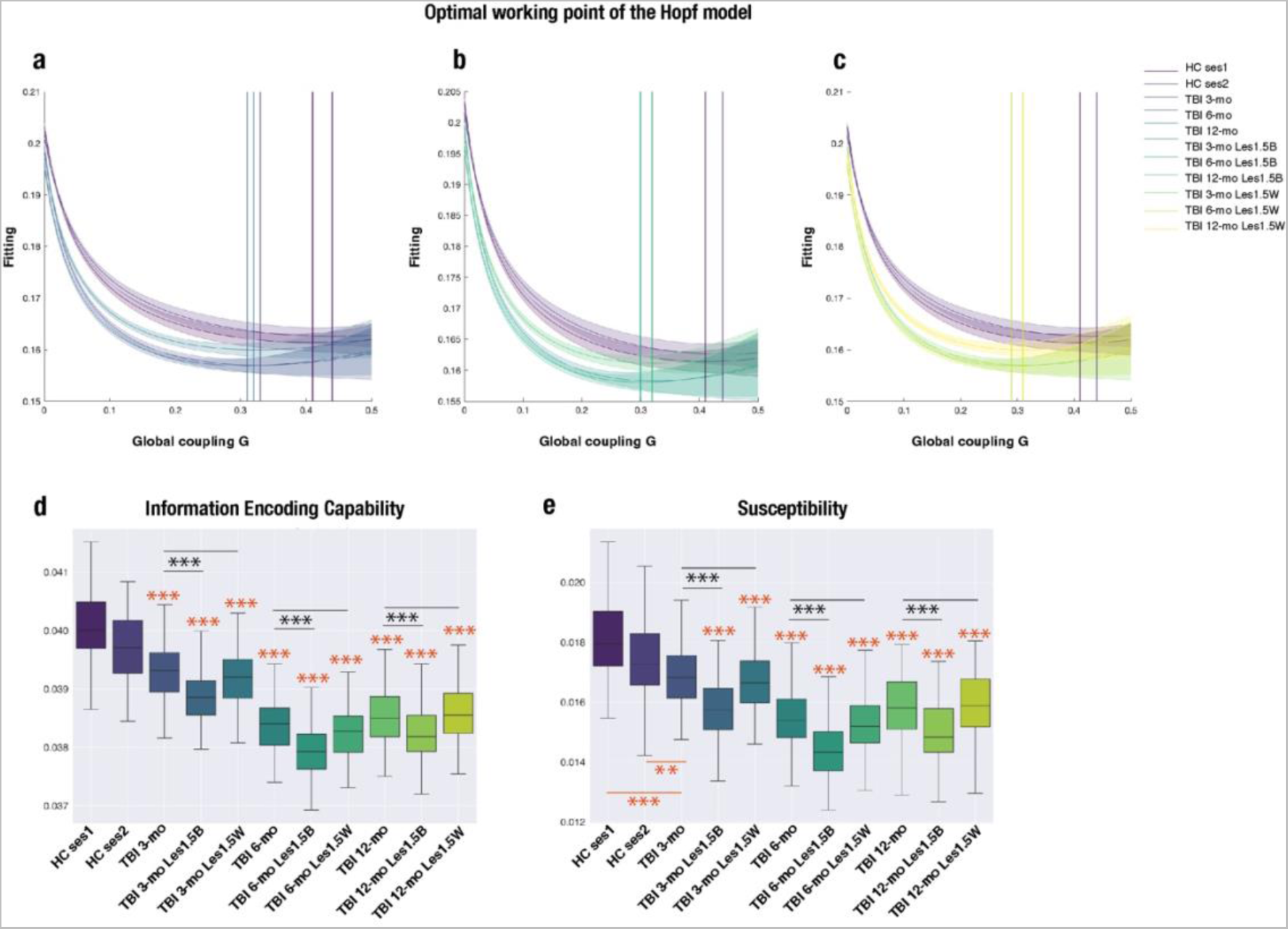
TBI patients show reduced reactivity to *in silico* perturbations over time as compared with healthy controls with the lowest value at 6-months post-injury with intact structural connectivity (SC) and after the simulated attack. **(a-c)** Evolution of the error of fitting the whole-brain model to the rs-fMRI data as a function of the global coupling strength, G. The optimal working point fitting the TBI data was lower than that for HCs in all conditions: with intact SC or after a simulated attack with binary and weighted lesion masks thresholded at 1.5 SD from the group’s mean volume of overlap with the Schaefer 1,000 nodes parcellation (see Material and methods). The shadow indicates the SEM. **(d)** Information encoding capability and **(e)** susceptibility measures were significantly different between HCs and TBI patients across all three time points with the lowest value shown by TBI patients at 6-months post-injury in all conditions. Further, the values with the binary approach for the simulated attack were also lower than with the weighted approach. The lower and upper edges of the box indicate the first (25th percentile) and third quartile (75th percentile), middle lines are medians. Orange asterisks denote comparisons between HCs session 1 and 2 and TBI patients (*** p < 0.0001, ** p < 0.01, results after Wilcoxon rank sum test comparing the distribution of values in 100 trials per condition). Black asterisks denote comparisons between intact SC and simulated attack approaches (*** p < 0.001, results after Wilcoxon signed rank test comparing the distribution of values in 100 trials per condition). Abbreviations: mo: months, les: lesion, 1.5: 1.5SD threshold, B: binary, W: weighted.

## 4. Discussion

Here we investigated how TBI impacts whole-brain resting state fMRI dynamics using a turbulence-based framework (Deco & Kringelbach, 2020; Deco et al., 2021). Our findings revealed specific spatial patterns of reduction in global amplitude turbulence between HCs and TBI patients at all three time points (3-, 6- and 12-months post-injury) differentially affecting long distances in the brain. Noteworthily, the greatest mean difference was observed between HCs and TBI patients at 6-months at all significant λ values. This was also the stage of recovery where we found differences in amplitude turbulence over time at long distances. Between group differences were also found at long distances for the other turbulent measures except for information transfer, where short distances were affected and no differences found in the longitudinal analysis. Importantly, when analyzing network-level turbulence between HCs and TBI patients for λ=0.03 (long distances), we found that, after a significant decrease at 6-months post-injury, amplitude turbulence recovers healthy levels in all RSNs except for the limbic by 12-months. Finally, the results from the whole-brain model indicated that susceptibility and information encoding capability are significantly reduced after TBI, with the largest difference again in the comparison between HCs and TBI patients at 6-months post-injury regardless of whether an intact or lesioned SC was used as input to the model. These results provide causal links needed for a better understanding of the recovery process one year after TBI. To the best of our knowledge, this is the first study to examine the longitudinal evolution of turbulent dynamics during recovery from TBI.

Our empirical finding of reduced turbulence at long distances after TBI as compared to HCs is in alignment with previous work showing a reduction in global metastability in TBI patients as compared with healthy controls (Hellyer et al., 2015). Furthermore, the turbulent dynamics framework adopted here lends indirect support to the graph theory results obtained in the same dataset (Roy et al., 2017) and extends those findings by offering a dynamical approach across spatial scales and a causal perspective on the information processing deficits following TBI during one year. Previously, Roy et al. (2017) have shown that global functional hyperconnectivity, in terms of network strength, peaked at 6-months post-injury. Of note, the local analysis revealed hyperconnectivity in the left frontal DMN and temporo-parietal attentional control networks across time points but was only associated with an increased cost at 6-months post-injury. Moreover, a cost-efficiency analysis showed that this elevated cost was mostly due to a higher number of medium-range connections rather than long-range connections. This fits well with our network-level finding where turbulence is reduced at 6-months post-injury in most RSNs with no residual effects after 12-months, when there was no cost associated with hyperconnectivity. Remarkably, we also found that the 15% quantile nodes in the HCs vs. TBI patients at 6-months were in the SM, DAT and CNT networks, in accordance with increased cost found in the temporo-parietal control network by Roy et al. (2017). Overall, our network-level findings suggest a shift in the functional plasticity of the brain in response to TBI where the turbulent dynamics at large distances rebalances to a healthy regime during the period spanning from 6 months to 12 months post-injury.

Regarding brain-behavior relationships, we found that performance on the Digit Span task (Forward and Backward) was positively associated with information cascade flow and amplitude turbulence in the DMN at long distances. This task is used to evaluate working memory processes, which are commonly affected after TBI, and requires the rapid communication across large-scale brain networks involving sensory (patients are read a sequence of numbers) and higher-order regions responsible for the manipulation of information for temporary use. Our findings suggest that the propagation of information between consecutive spatial scales and higher levels of amplitude in the DMN at rest might be an important dynamical mechanism to support working memory tasks. However, future work with larger sample sizes could better characterize additional associations between turbulent measures and performance in other cognitive domains as this has been previously found for metastability and cognitive flexibility, information processing and associative memory after TBI (Hellyer et al., 2015).

Our computational findings are consistent with our empirical results, demonstrating that TBI hinders the reactivity of the brain to external *in silico* perturbations. When fitting the Hopf model to each empirical condition, we found lower values in the optimal working point for TBI patients across time points regardless of whether an intact or lesioned SC was used as input to the model. However, significant differences between the binary and the weighted approach emerged when examining the susceptibility and information encoding capability after our *in silico* perturbation. Indeed, the values of these measures for the binary simulated attack were significantly reduced at all stages of recovery with respect to the values in the weighted simulated attack. This is indicative that these measures are less affected by a distributed pattern of lesions, as the one in this cohort of TBI patients, and may be more sensitive to focal lesions that greatly overlap in the sample. Whether this result is contingent upon the anatomical identity (and connectivity profile) of the lesions remains an interesting question for future research. That said, we believe that this is an interesting finding on its own, highlighting the importance of the impact that focal lesions may have in other patients’ populations, such as stroke, where usually most patients present a lesion in the vascular territory of the middle cerebral artery and there is greater degree of overlap.

The results from the present study have some potential clinical implications. Turbulent measures might be used to improve clinical diagnosis and prognosis after TBI as they demonstrated sensitivity to the longitudinal changes in whole-brain dynamics compared to HCs. However, further information regarding the relationship of these measures and TBI severity is still needed and calls for collaborative efforts to perform these analyses on large cohorts of patients. On the other hand, information processing-based measures obtained from resting state data could provide a meaningful and repeatable way to monitor the evolution of TBI, the recovery progress, and the patient’s possible response to treatments as these remain prominent challenges in clinical settings to date (National Academies of Sciences et al., 2022). Our *in silico* perturbation approach with the Hopf computational model also indicated reduced coupling and reactivity to external stimulation after TBI across time points. We restricted our perturbative protocol to the introduction of random values in all nodes to modify the local bifurcation parameter of the model. That being said, it would be interesting to implement an excitatory protocol for individual nodes or a subset of relevant nodes in order to identify brain targets that can be stimulated with techniques such as transcranial magnetic stimulation (TMS) (Demirtas-Tatlidede et al., 2012). Notably, some promising results indicate that TMS and neurorehabilitation may act synergistically to improve functional outcomes after TBI (Nardone et al., 2020). Yet, we leave the implementation of such perturbative protocol for future work.

It is also worthwhile to acknowledge some limitations in the current study. Although this dataset represents a relatively homogenous sample in terms of age and time since injury, it remains of small size and a replication with a larger cohort of patients is needed to assess the generalizability of these results. In addition, the resting state sequence comprised 150 scans with a temporal resolution of 2s that were acquired over a 5-minute period. While it has been shown that functional correlation of RSNs only stabilizes after 5-min with TR=2.3s (Van Dijk et al., 2010), recent evidence indicates that metastability is less affected by temporal resolution and shows greater reliability in short-term scans (Yang et al., 2021). Even if our simulated attack approach allowed us to take into account the effect of focal lesions on the SC, it leaves aside inter-individual differences in SC and future studies with personalized models are warranted to validate our findings. Finally, although we were able to partially replicate our results using the 400 node Schaefer parcellation, it remains unknown how they may be affected when using a different atlas.

In conclusion, we have shown that whole-brain turbulent dynamics complement previous findings on the temporal evolution of functional connectivity after TBI in an open access dataset of healthy controls and patients with data collected after 3-, 6- and 12-months post-injury. Specifically, our empirical results suggest the presence of maladaptive neuroplasticity at 6-months post-injury as manifested in the reduction of global amplitude turbulence at long distances. Thus, we demonstrated that the framework of turbulent dynamics may facilitate our understanding of how the brain responds to traumatic injury. Importantly, our computational approach suggests that in this cohort of moderate-to-severe TBI patients, their brain is the least reactive to external perturbations at 6-months post-injury during the recovery trajectory.

## CRediT author statement

**Noelia Martínez-Molina:** Conceptualization, Data curation, Formal analysis, Visualization, Writing - original draft, Project administration. **Anira Escrichs:** Software and Writing - review & editing. **Yonatan Sanz-Perl:** Software, Validation and Writing - review & editing. **Aleksi J. Sihvonen:** Writing - review & editing. **Teppo Särkämö:** Writing - review & editing. **Morten L. Kringelbach:** Methodology, Visualization, Software and Writing - review & editing. **Gustavo Deco:** Supervision, Methodology, Software and Writing - review & editing.

## Code availability statement

All code written in support of this study is publicly available on https://github.com/noechan/TBI_ON_turbu_Hopf_NC

## Technical terms

**Diffuse axonal injury:** damage caused to axons and axoplasmic membranes by shearing forces produced at the time of brain injury. This impairs axonal transport and disrupts brain function and structure.

**Turbulence:** In coupled oscillators, turbulence is characterized by the high variability across space and time of the Kuramoto local order parameter capturing the local synchronization, so in this sense it is a spatiotemporal extension of the level of metastability.

**Information transfer:** A measure that characterizes how the information propagates across space at a particular scale.

**Information cascade flow:** A measure that estimates the stream of information between a given scale and a subsequent lower scale in consecutive time steps.

**Information cascade:** A measure that captures the entire efficient information-processing behavior across scales.

**Metastability:** Fundamental concept describing complex systems’ behavior in terms of the variability in the global state of synchronization as a function of time.

## Supporting Information

### Outlier identification and denoising in the CONN toolbox

Outlier scans were identified as those that exceeded 3 standard deviations in z-scores from the global BOLD signal change (GSC) or with framewise displacement (FD) that exceeded 2 mm. The GSC timeseries was computed at each scan as the absolute value of the scan-to-scan change in global BOLD signal using SPM global BOLD signal definition. The GSC timeseries were then scaled to standard units within each run by subtracting their median value and dividing by 0.74 times their interquartile range (Whitfield-Gabrieli et al., 2011). The FD timeseries was defined as the maximum change in the position of six points placed at the centers of each face in a 140 x 180 x 115 mm bounding box around the brain and undergoing the same rotations and translations as the participant’s head.

For denoising the timeseries, we used the default pipeline in the CONN toolbox (Nieto-Castanon, 2020) in order to characterize (noise components extraction) and minimize the effect of non-neural noise on the BOLD timeseries. Participant-specific minimally eroded white matter (WM) and cerebrospinal fluid (CSF) masks were generated through a one-voxel binary erosion of tissue masks obtained after segmentation. Next, noise components were extracted from these WM and CSF masks using principal components analysis following the anatomical aCompCor methods (Behzadi et al., 2007). Principal component from WM and CSF were computed after discounting motion and outlier effects. Ordinary least square regression removed from each voxel BOLD timeseries the effect of all identified noise components including 5 components from WM, 5 components from CSF, 12 estimated motion parameters (6 realignment parameters and their first order time derivatives), outlier scans, effect of session and its first level derivative convolved with the canonical hemodynamic response function, and constant and linear session effects (i.e., linear detrending).

### Quality Control in the CONN toolbox

Participant-level visual quality control (QC) was done to evaluate possible patterns or other features that may be visible in the BOLD signal timeseries after denoising. We reviewed run-specific plots rendering carpetplots (Power, 2017) of fully preprocessed BOLD timeseries before and after denoising, together with the traces of GSC, FD and outliers timeseries. This inspection aimed to confirm that sudden and synchronized variations in signal intensity had been flagged as outliers, and that there were no visible residual large-scale patterns in the BOLD signal timeseries, which could indicate the persistence of global or widespread noise sources.

In addition, we computed functional connectivity values (Pearson’s correlation coefficients) among all pairs from a fixed set of 1,000 random voxels within the MNI-space gray matter template mask in order to evaluate a relatively dense sample of connections from the whole-brain connectome. From these connections, we computed and displayed the distribution of functional connectivity (FC) values separately for each participant’s functional run. Visual inspection of these distributions allows us to evaluate the relative presence of residual noise sources in the BOLD timeseries, which tend to shift the entire FC distribution towards positive values (false positives), altering the FC distribution center and overall shape in a manner that is highly variable across runs. In comparison, the relative absence of noise sources is expressed as FC distributions that appear relatively centered (with a small positive distribution mean and a distribution mode approximately at zero) and similar across different runs and participants (Nieto-Castanon 2020). After evaluating the distribution of functional connectivity after denoising, the number of noise components extracted from the WM was increased to 10.

**Table S1.**
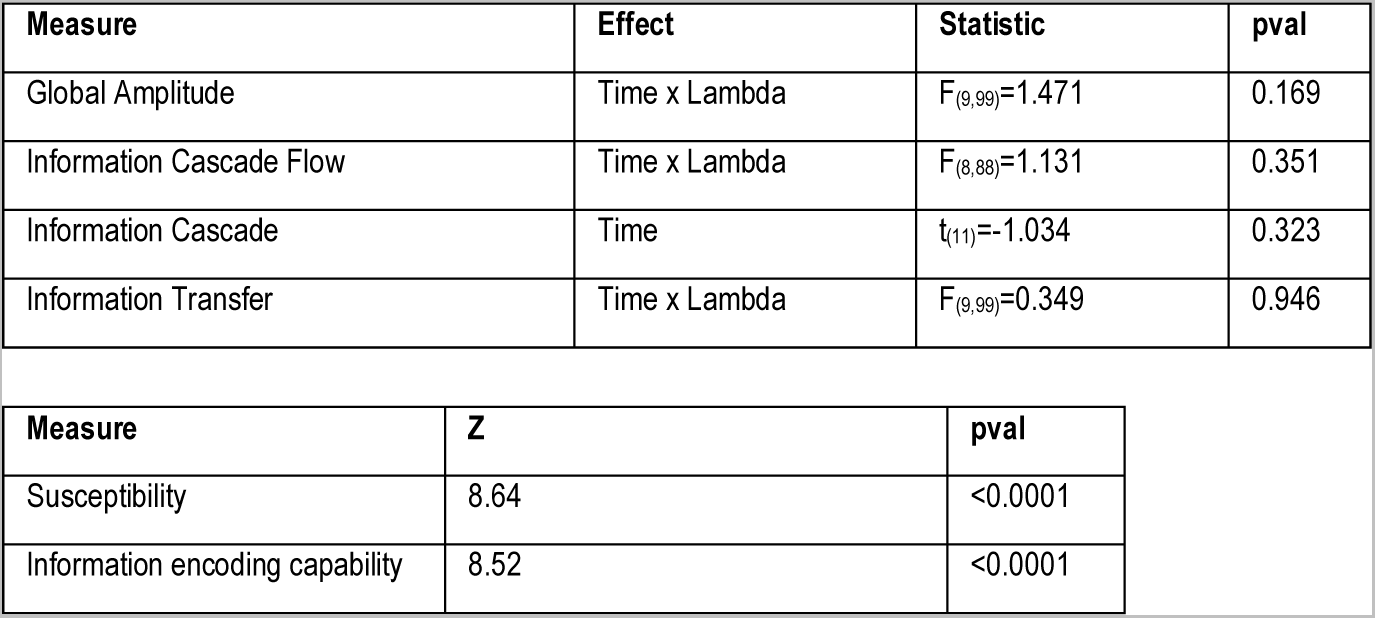
Turbulent and perturbational measures in Healthy Controls. Results from the repeated measures ANOVA with time as within-group factor and paired T-test (top table) and Wilcoxon signed rank test (bottom table). <u>Schaefer parcellation 1000 nodes 7 RSNs</u>.

| Measure | Effect | Statistic | pval |
| --- | --- | --- | --- |
| Global Amplitude | Time x Lambda | $F_{(9,99)}=1.471$ | 0.169 |
| Information Cascade Flow | Time x Lambda | $F_{(8,88)}=1.131$ | 0.351 |
| Information Cascade | Time | $t_{(11)}=-1.034$ | 0.323 |
| Information Transfer | Time x Lambda | $F_{(9,99)}=0.349$ | 0.946 |

| Measure | Z | pval |
| --- | --- | --- |
| Susceptibility | 8.64 | <0.0001 |
| Information encoding capability | 8.52 | <0.0001 |

**Table S2.** Global Amplitude Turbulence in Healthy Controls (mean across session 1 and 2) vs TBI patients (3-,6- and 12-months). Results from the 4 (Group) x 10 (Lambda) ANOVA (top table) and post hoc tests (bottom table). <u>Schaefer parcellation 1000 nodes 7 RSNs</u>.

| Effect | $F_{(27,440)}$ | p-val | Effect size ($\eta^2_{\text{partial}}$) |
| --- | --- | --- | --- |
| Group x Lambda | 2.534 | <0.0001 | 0.135 |

| Lambda, Group | Lambda, Group | Mean Difference | t | pval |
| --- | --- | --- | --- | --- |
| 0.01, Healthy Controls | 0.01, TBI at 3-months post-injury | 0.014 | 4.201 | **0.018** |
|  | 0.01, TBI at 6-months post-injury | **0.024** | 7.296 | **<0.0001** |
|  | 0.01, TBI at 12-months post-injury | 0.014 | 4.208 | **0.017** |
| 0.03, Healthy Controls | 0.03, TBI at 3-months post-injury | 0.015 | 4.760 | **0.002** |
|  | 0.03, TBI at 6-months post-injury | **0.023** | 7.108 | **<0.0001** |
|  | 0.03, TBI at 12-months post-injury | 0.014 | 4.152 | **0.022** |
| 0.06, Healthy Controls | 0.06, TBI at 3-months post-injury | 0.014 | 4.166 | **0.020** |
|  | 0.06, TBI at 6-months post-injury | **0.017** | 5.326 | **<0.0001** |
|  | 0.06, TBI at 12-months post-injury | 0.009 | 2.898 | 0.607 |
| 0.09, Healthy Controls | 0.09, TBI at 3-months post-injury | 0.010 | 3.166 | 0.386 |
|  | 0.09, TBI at 6-months post-injury | 0.012 | 3.676 | 0.108 |
|  | 0.09, TBI at 12-months post-injury | 0.006 | 1.925 | 0.998 |
| 0.12, Healthy Controls | 0.12, TBI at 3-months post-injury | 0.007 | 2.271 | 0.964 |
|  | 0.12, TBI at 6-months post-injury | 0.008 | 2.467 | 0.901 |
|  | 0.12, TBI at 12-months post-injury | 0.004 | 1.297 | 1.000 |
| 0.15, Healthy Controls | 0.15, TBI at 3-months post-injury | 0.005 | 1.592 | 1.000 |
|  | 0.15, TBI at 6-months post-injury | 0.005 | 1.641 | 1.000 |
|  | 0.15, TBI at 12-months post-injury | 0.003 | 1.015 | 1.000 |
| 0.18, Healthy Controls | 0.18, TBI at 3-months post-injury | 0.004 | 1.151 | 1.000 |
|  | 0.18, TBI at 6-months post-injury | 0.004 | 1.107 | 1.000 |
|  | 0.18, TBI at 12-months post-injury | 0.003 | 0.888 | 1.000 |
| 0.21, Healthy Controls | 0.21, TBI at 3-months post-injury | 0.003 | 0.871 | 1.000 |
|  | 0.21, TBI at 6-months post-injury | 0.002 | 0.754 | 1.000 |
|  | 0.21, TBI at 12-months post-injury | 0.003 | 0.793 | 1.000 |
| 0.24, Healthy Controls | 0.24, TBI at 3-months post-injury | 0.002 | 0.671 | 1.000 |
|  | 0.24, TBI at 6-months post-injury | 0.002 | 0.503 | 1.000 |
|  | 0.24, TBI at 12-months post-injury | 0.002 | 0.696 | 1.000 |
| 0.27, Healthy Controls | 0.27, TBI at 3-months post-injury | 0.002 | 0.521 | 1.000 |
|  | 0.27, TBI at 6-months post-injury | 0.001 | 0.324 | 1.000 |
|  | 0.27, TBI at 12-months post-injury | 0.002 | 0.602 | 1.000 |

**Table S3.** Information Cascade Flow in Healthy Controls (mean across session 1 and 2) vs TBI patients (3-,6- and 12-months). Results from the 4 (Group) x 9 (Lambda) ANOVA (top table) and post hoc tests (bottom table). <u>Schaefer parcellation 1000 nodes 7 RSNs</u>.

| Effect | F<sub>(24,396)</sub> | p-val | Effect size ($\eta^2_{\text{partial}}$) |
| --- | --- | --- | --- |
| Group x Lambda | 3.007 | <0.0001 | 0.154 |

| Lambda, Group | Lambda, Group | Mean Difference | t | pval |
| --- | --- | --- | --- | --- |
| 0.01, Healthy Controls | 0.01, TBI at 3-months post-injury | 0.055 | 4.512 | **0.004** |
|  | 0.01, TBI at 6-months post-injury | 0.106 | 8.718 | **0.000** |
|  | 0.01, TBI at 12-months post-injury | 0.047 | 3.873 | **0.049** |
| 0.03, Healthy Controls | 0.03, TBI at 3-months post-injury | 0.038 | 3.098 | 0.394 |
|  | 0.03, TBI at 6-months post-injury | 0.064 | 5.303 | **<0.0001** |
|  | 0.03, TBI at 12-months post-injury | 0.033 | 2.700 | 0.716 |
| 0.06, Healthy Controls | 0.06, TBI at 3-months post-injury | 0.020 | 1.619 | 1.000 |
|  | 0.06, TBI at 6-months post-injury | 0.029 | 2.356 | 0.918 |
|  | 0.06, TBI at 12-months post-injury | 0.016 | 1.342 | 1.000 |
| 0.09, Healthy Controls | 0.09, TBI at 3-months post-injury | 0.013 | 1.040 | 1.000 |
|  | 0.09, TBI at 6-months post-injury | 0.015 | 1.273 | 1.000 |
|  | 0.09, TBI at 12-months post-injury | 0.010 | 0.812 | 1.000 |
| 0.12, Healthy Controls | 0.12, TBI at 3-months post-injury | 0.008 | 0.695 | 1.000 |
|  | 0.12, TBI at 6-months post-injury | 0.009 | 0.751 | 1.000 |
|  | 0.12, TBI at 12-months post-injury | 0.006 | 0.473 | 1.000 |
| 0.15, Healthy Controls | 0.15, TBI at 3-months post-injury | 0.006 | 0.461 | 1.000 |
|  | 0.15, TBI at 6-months post-injury | 0.006 | 0.479 | 1.000 |
|  | 0.15, TBI at 12-months post-injury | 0.003 | 0.266 | 1.000 |
| 0.18, Healthy Controls | 0.18, TBI at 3-months post-injury | 0.004 | 0.312 | 1.000 |
|  | 0.18, TBI at 6-months post-injury | 0.004 | 0.322 | 1.000 |
|  | 0.18, TBI at 12-months post-injury | 0.002 | 0.138 | 1.000 |
| 0.21, Healthy Controls | 0.21, TBI at 3-months post-injury | 0.003 | 0.219 | 1.000 |
|  | 0.21, TBI at 6-months post-injury | 0.003 | 0.229 | 1.000 |
|  | 0.21, TBI at 12-months post-injury | 7.924×10<sup>-4</sup> | 0.065 | 1.000 |
| 0.24, Healthy Controls | 0.24, TBI at 3-months post-injury | 0.002 | 0.167 | 1.000 |
|  | 0.24, TBI at 6-months post-injury | 0.002 | 0.169 | 1.000 |
|  | 0.24, TBI at 12-months post-injury | 3.200×10<sup>-4</sup> | 0.026 | 1.000 |

**Table S4.**
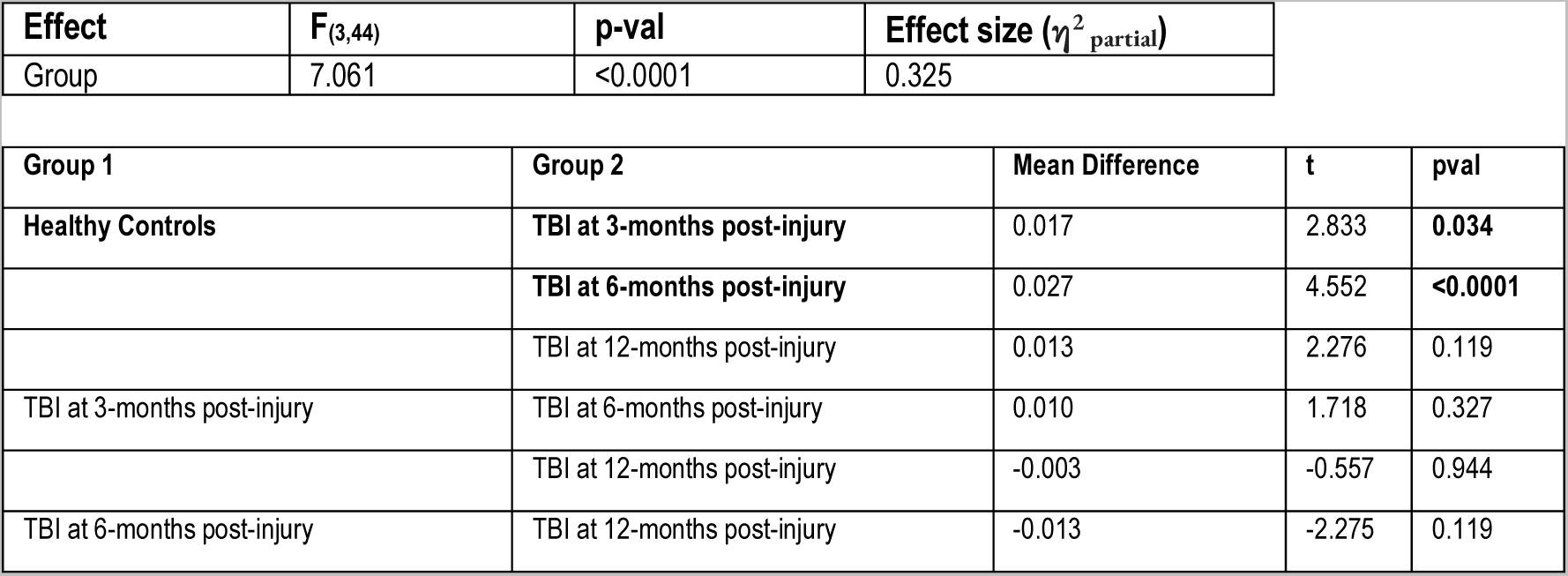
Information Cascade in Healthy Controls (mean across session 1 and 2) vs TBI patients (3-,6- and 12-months). Results from the 4 one-way ANOVA (top table) and post hoc tests (bottom table). <u>Schaefer</u> <u>parcellation 1000 nodes 7 RSNs</u>.

**Table S5.** Information Transfer in Healthy Controls Healthy Controls (mean across session 1 and 2) vs TBI patients (3-,6- and 12-months). Results from the 4 (Group) x 10 (Lambda) ANOVA (top table) and post hoc tests (bottom table). <u>Schaefer parcellation 1000 nodes 7 RSNs</u>.

| Effect | F<sub>(27,440)</sub> | p-val | Effect size ($\eta^2_{\text{partial}}$) |
| --- | --- | --- | --- |
| Group x Lambda | 2.325 | <0.0001 | 0.125 |

| Lambda, Group | Lambda, Group | Mean Difference | t | pval |
| --- | --- | --- | --- | --- |
| 0.01, Healthy Controls | 0.01, TBI at 3-months post-injury | -4.843×10<sup>-4</sup> | -0.023 | 1.000 |
|  | 0.01, TBI at 6-months post-injury | -0.001 | -0.057 | 1.000 |
|  | 0.01, TBI at 12-months post-injury | -6.715×10<sup>-4</sup> | -0.032 | 1.000 |
| 0.03, Healthy Controls | 0.03, TBI at 3-months post-injury | -0.007 | -0.321 | 1.000 |
|  | 0.03, TBI at 6-months post-injury | -0.011 | -0.514 | 1.000 |
|  | 0.03, TBI at 12-months post-injury | -0.007 | -0.324 | 1.000 |
| 0.06, Healthy Controls | 0.06, TBI at 3-months post-injury | -0.026 | -1.225 | 1.000 |
|  | 0.06, TBI at 6-months post-injury | -0.028 | -1.321 | 1.000 |
|  | 0.06, TBI at 12-months post-injury | -0.020 | -0.965 | 1.000 |
| 0.09, Healthy Controls | 0.09, TBI at 3-months post-injury | -0.048 | -2.313 | 0.954 |
|  | 0.09, TBI at 6-months post-injury | -0.042 | -2.014 | 0.994 |
|  | 0.09, TBI at 12-months post-injury | -0.033 | -1.605 | 1.000 |
| 0.12, Healthy Controls | 0.12, TBI at 3-months post-injury | -0.073 | -3.500 | 0.177 |
|  | 0.12, TBI at 6-months post-injury | -0.055 | -2.627 | 0.813 |
|  | 0.12, TBI at 12-months post-injury | -0.044 | -2.110 | 0.988 |
| **0.15, Healthy Controls** | **0.15, TBI at 3-months post-injury** | -0.095 | -4.559 | **0.004** |
|  | 0.15, TBI at 6-months post-injury | -0.067 | -3.199 | 0.362 |
|  | 0.15, TBI at 12-months post-injury | -0.054 | -2.590 | 0.837 |
| **0.18, Healthy Controls** | **0.18, TBI at 3-months post-injury** | -0.112 | -5.389 | **<0.0001** |
|  | 0.18, TBI at 6-months post-injury | -0.077 | -3.681 | 0.106 |
|  | 0.18, TBI at 12-months post-injury | -0.064 | -3.088 | 0.449 |
| **0.21, Healthy Controls** | **0.21, TBI at 3-months post-injury** | -0.126 | -6.046 | **<0.0001** |
|  | **0.21, TBI at 6-months post-injury** | -0.085 | -4.062 | **0.030** |
|  | 0.21, TBI at 12-months post-injury | -0.074 | -3.558 | 0.151 |
| **0.24, Healthy Controls** | **0.24, TBI at 3-months post-injury** | -0.138 | -6.631 | **<0.0001** |
|  | **0.24, TBI at 6-months post-injury** | -0.092 | -4.416 | **0.008** |
|  | **0.24, TBI at 12-months post-injury** | -0.084 | -4.025 | **0.034** |
| **0.27, Healthy Controls** | **0.27, TBI at 3-months post-injury** | -0.143 | -6.848 | **<0.0001** |
|  | **0.27, TBI at 6-months post-injury** | -0.099 | -4.755 | **0.002** |
|  | **0.27, TBI at 12-months post-injury** | -0.093 | -4.474 | **0.006** |

**Table S6.** Global Amplitude Turbulence in Healthy Controls (mean across session 1 and 2) vs TBI patients (3-,6- and 12-months). Results from the 4 (Group) x 10 (Lambda) ANOVA (top table) and post hoc tests (bottom table). <u>Schaefer parcellation 400 nodes 7 RSNs</u>.

| **Effect** | **F<sub>(27,440)</sub>** | **p-val** | **Effect size ($\eta^2_{\text{partial}}$)** |
| --- | --- | --- | --- |
| Group x Lambda | 3.179 | <0.0001 | 0.163 |

| **Lambda, Group** | **Lambda, Group** | **Mean Difference** | **t** | **pval** |
| --- | --- | --- | --- | --- |
| **0.01, Healthy Controls** | **0.01, TBI at 3-months post-injury** | 0.016 | 4.635 | **0.003** |
|  | **0.01, TBI at 6-months post-injury** | 0.026 | 7.410 | **<0.0001** |
|  | **0.01, TBI at 12-months post-injury** | 0.018 | 5.218 | **<0.0001** |
| **0.03, Healthy Controls** | 0.03, TBI at 3-months post-injury | 0.012 | 3.605 | 0.132 |
|  | **0.03, TBI at 6-months post-injury** | 0.020 | 5.913 | **<0.0001** |
|  | 0.03, TBI at 12-months post-injury | 0.013 | 3.637 | 0.121 |
| 0.06, Healthy Controls | 0.06, TBI at 3-months post-injury | 0.003 | 0.931 | 1.000 |
|  | 0.06, TBI at 6-months post-injury | 0.008 | 2.218 | 0.974 |
|  | 0.06, TBI at 12-months post-injury | 0.001 | 0.425 | 1.000 |
| 0.09, Healthy Controls | 0.09, TBI at 3-months post-injury | 0.003 | 0.919 | 1.000 |
|  | 0.09, TBI at 6-months post-injury | 0.005 | 1.438 | 1.000 |
|  | 0.09, TBI at 12-months post-injury | 0.001 | 0.308 | 1.000 |
| 0.12, Healthy Controls | 0.12, TBI at 3-months post-injury | 0.012 | 3.574 | 0.145 |
|  | 0.12, TBI at 6-months post-injury | 0.013 | 3.747 | 0.087 |
|  | 0.12, TBI at 12-months post-injury | 0.011 | 3.191 | 0.367 |
| **0.15, Healthy Controls** | **0.15, TBI at 3-months post-injury** | 0.021 | 5.994 | **<0.0001** |
|  | **0.15, TBI at 6-months post-injury** | 0.021 | 6.035 | **<0.0001** |
|  | **0.15, TBI at 12-months post-injury** | 0.020 | 5.945 | **<0.0001** |
| **0.18, Healthy Controls** | **0.18, TBI at 3-months post-injury** | 0.024 | 7.050 | **<0.0001** |
|  | **0.18, TBI at 6-months post-injury** | 0.024 | 7.042 | **<0.0001** |
|  | **0.18, TBI at 12-months post-injury** | 0.025 | 7.189 | **<0.0001** |
| **0.21, Healthy Controls** | **0.21, TBI at 3-months post-injury** | 0.025 | 7.113 | **<0.0001** |
|  | **0.21, TBI at 6-months post-injury** | 0.024 | 7.094 | **<0.0001** |
|  | 0.21, TBI at 12-months post-injury | 0.025 | 7.336 | <0.0001 |
| 0.24, Healthy Controls | 0.24, TBI at 3-months post-injury | 0.023 | 6.571 | <0.0001 |
|  | 0.24, TBI at 6-months post-injury | 0.023 | 6.558 | <0.0001 |
|  | 0.24, TBI at 12-months post-injury | 0.023 | 6.809 | <0.0001 |
| 0.27, Healthy Controls | 0.27, TBI at 3-months post-injury | 0.020 | 5.737 | <0.0001 |
|  | 0.27, TBI at 6-months post-injury | 0.020 | 5.732 | <0.0001 |
|  | 0.27, TBI at 12-months post-injury | 0.021 | 5.955 | <0.0001 |

**Table S7.**
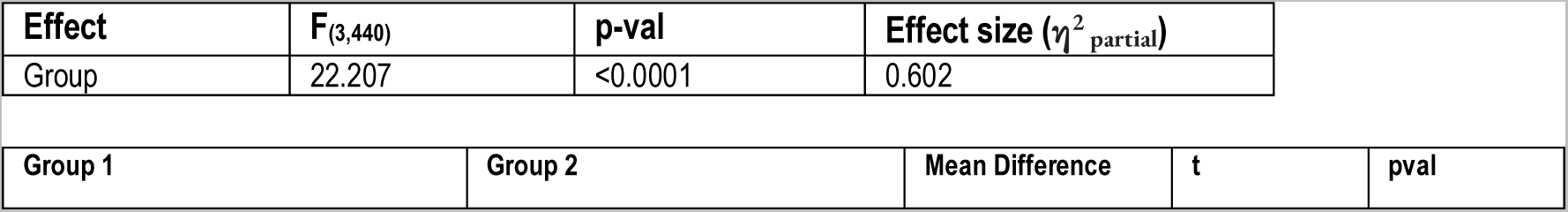
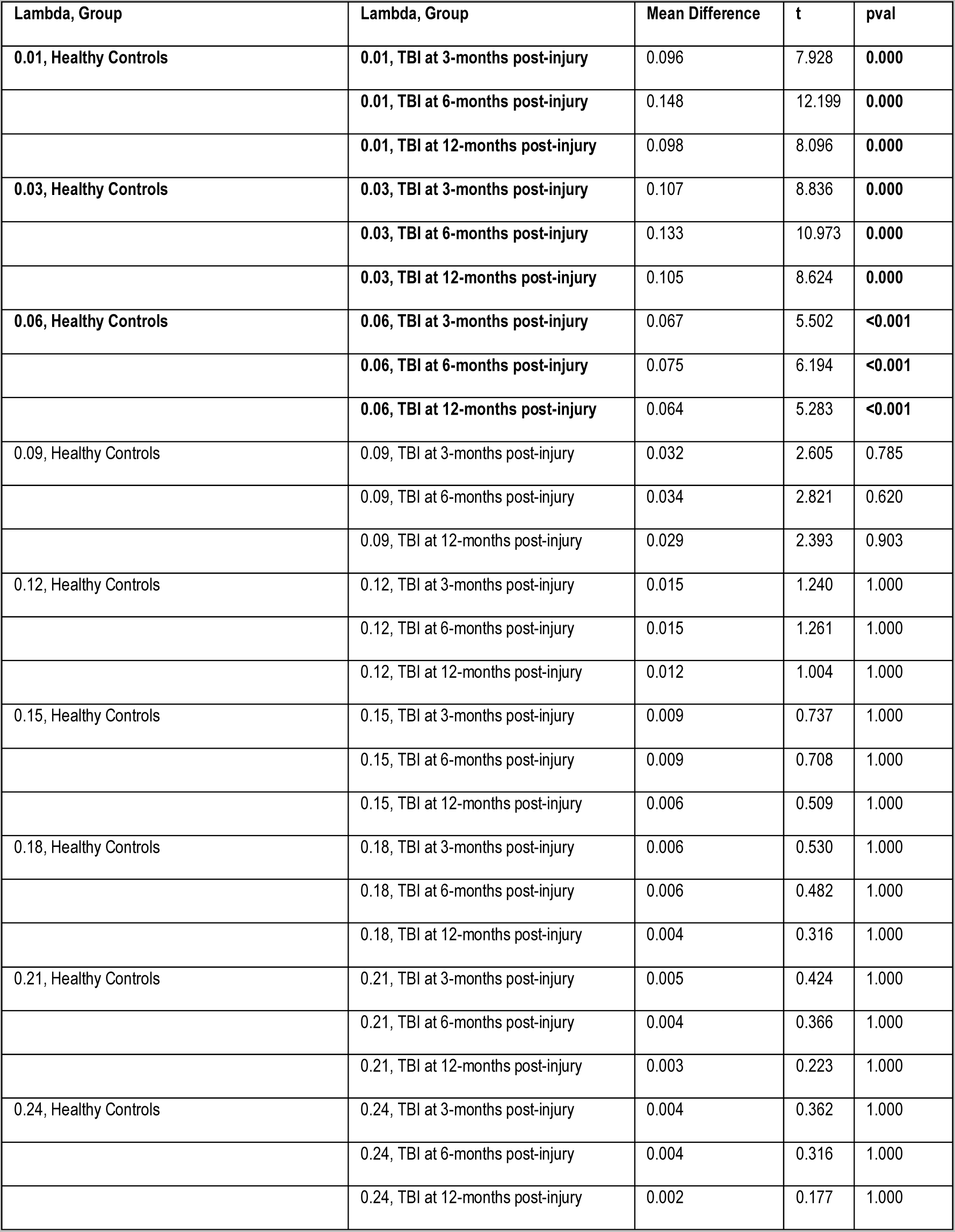
Information Cascade Flow in Healthy Controls (mean across session 1 and 2) vs TBI patients (3-,6- and 12-months). Results from the 4 (Group) x 9 (Lambda) ANOVA (top table) and post hoc tests (bottom table). <u>Schaefer parcellation 400 nodes 7 RSNs</u>.

**Table S8.**
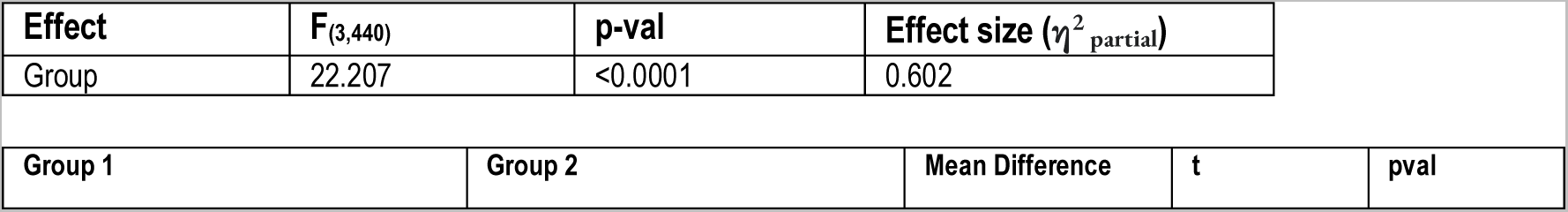
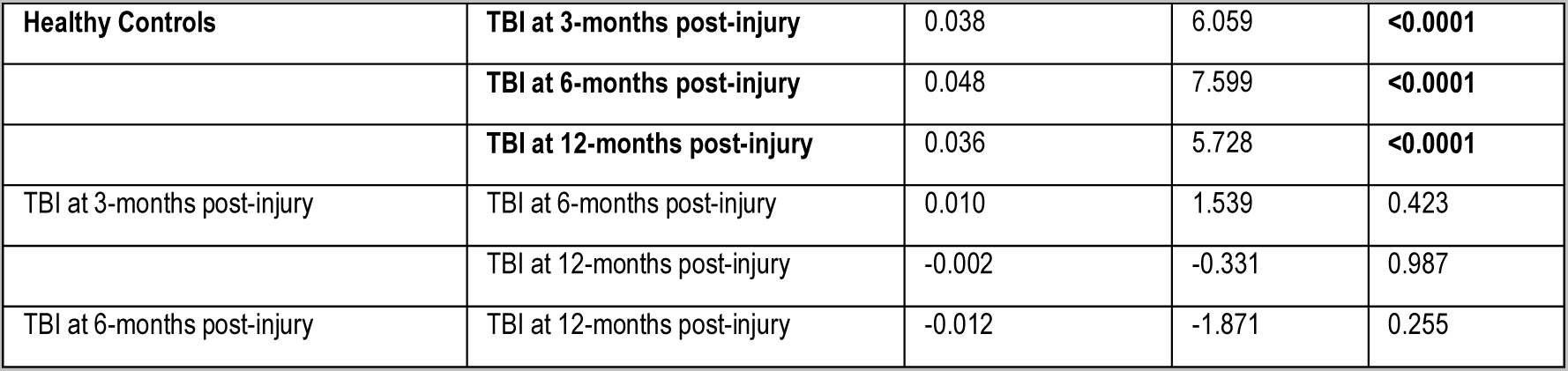
Information Cascade in Healthy Controls (mean across session 1 and 2) vs TBI patients (3-,6- and 12-months). Results from the 4 one-way ANOVA (top table) and post hoc tests (bottom table). <u>Schaefer</u> <u>parcellation 400 nodes 7 RSNs</u>.

**Table S9.** Information Transfer in Healthy Controls mean across session 1 and 2) vs TBI patients (3-,6- and 12-months). Results from the 4 (Group) x 10 (Lambda) ANOVA (top table) and post hoc tests (bottom table). <u>Schaefer parcellation 400 nodes 7 RSNs</u>.

| Effect | F<sub>(27,440)</sub> | p-val | Effect size ($\eta^2_{\text{partial}}$) |
| --- | --- | --- | --- |
| Group x Lambda | 3.305 | <0.0001 | 0.169 |

| Lambda, Group | Lambda, Group | Mean Difference | t | pval |
| --- | --- | --- | --- | --- |
| 0.01, Healthy Controls | 0.01, TBI at 3-months post-injury | -0.001 | -0.082 | 1.000 |
|  | 0.01, TBI at 6-months post-injury | -0.002 | -0.122 | 1.000 |
|  | 0.01, TBI at 12-months post-injury | -0.002 | -0.100 | 1.000 |
| 0.03, Healthy Controls | 0.03, TBI at 3-months post-injury | -0.015 | -1.001 | 1.000 |
|  | 0.03, TBI at 6-months post-injury | -0.021 | -1.356 | 1.000 |
|  | 0.03, TBI at 12-months post-injury | -0.017 | -1.112 | 1.000 |
| 0.06, Healthy Controls | 0.06, TBI at 3-months post-injury | -0.051 | -3.305 | 0.288 |
|  | 0.06, TBI at 6-months post-injury | -0.060 | -3.915 | 0.050 |
|  | 0.06, TBI at 12-months post-injury | -0.052 | -3.342 | 0.263 |
| **0.09, Healthy Controls** | 0.09, TBI at 3-months post-injury | -0.059 | -3.801 | 0.073 |
|  | **0.09, TBI at 6-months post-injury** | -0.069 | -4.492 | **0.006** |
|  | 0.09, TBI at 12-months post-injury | -0.058 | -3.779 | 0.078 |
| 0.12, Healthy Controls | 0.12, TBI at 3-months post-injury | -0.026 | -1.691 | 1.000 |
|  | 0.12, TBI at 6-months post-injury | -0.037 | -2.422 | 0.919 |
|  | 0.12, TBI at 12-months post-injury | -0.024 | -1.583 | 1.000 |
| 0.15, Healthy Controls | 0.15, TBI at 3-months post-injury | 0.008 | 0.520 | 1.000 |
|  | 0.15, TBI at 6-months post-injury | -0.003 | -0.222 | 1.000 |
|  | 0.15, TBI at 12-months post-injury | 0.012 | 0.806 | 1.000 |
| 0.18, Healthy Controls | 0.18, TBI at 3-months post-injury | 0.026 | 1.652 | 1.000 |
|  | 0.18, TBI at 6-months post-injury | 0.015 | 0.949 | 1.000 |
|  | 0.18, TBI at 12-months post-injury | 0.033 | 2.141 | 0.985 |
| 0.21, Healthy Controls | 0.21, TBI at 3-months post-injury | 0.036 | 2.306 | 0.956 |
|  | 0.21, TBI at 6-months post-injury | 0.026 | 1.659 | 1.000 |
|  | 0.21, TBI at 12-months post-injury | 0.046 | 2.994 | 0.526 |
| 0.24, Healthy Controls | 0.24, TBI at 3-months post-injury | 0.041 | 2.650 | 0.798 |
|  | 0.24, TBI at 6-months post-injury | 0.032 | 2.068 | 0.991 |
|  | 0.24, TBI at 12-months post-injury | 0.054 | 3.516 | 0.169 |
| 0.27, Healthy Controls | 0.27, TBI at 3-months post-injury | 0.043 | 2.750 | 0.727 |
|  | 0.27, TBI at 6-months post-injury | 0.035 | 2.240 | 0.970 |
|  | 0.27, TBI at 12-months post-injury | 0.058 | 3.779 | 0.078 |

**Table S10.**
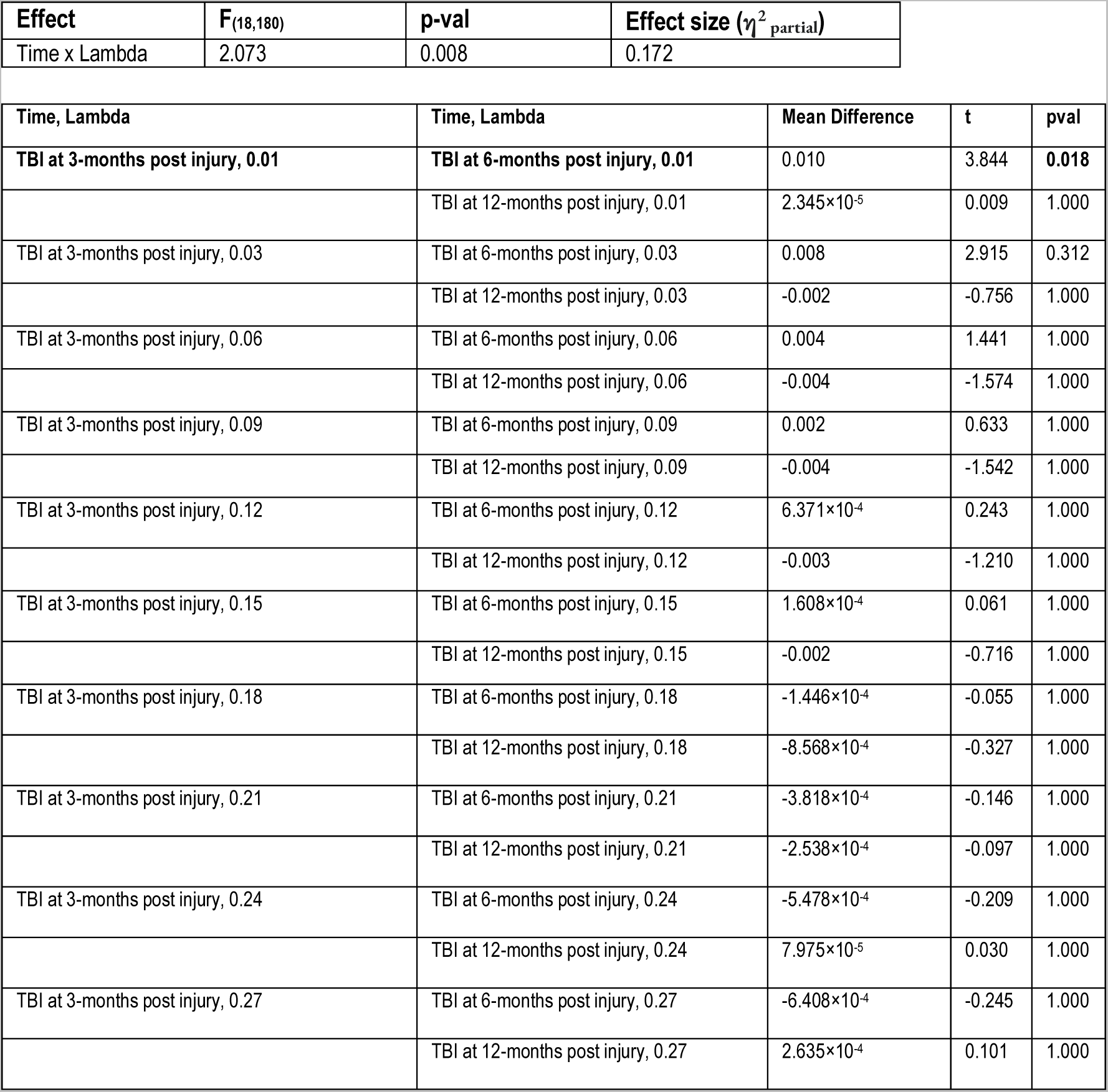
Global Amplitude Turbulence in TBI patients over time. Results from the repeated measures ANCOVA with lesion volume as nuisance covariate (top table) and post hoc tests (bottom table).

**Table S11.** Information Cascade Flow in TBI patients over time. Results from the repeated measures ANCOVA with lesion volume as nuisance covariate (top table) and post hoc tests (bottom table).

| Effect | F<sub>(16,160)</sub> | p-val | Effect size ($\eta^2_{\text{partial}}$) |
| --- | --- | --- | --- |
| Time x Lambda | 1.809 | 0.034 | 0.153 |

| Time, Lambda | Time, Lambda | Mean Difference | t | pval |
| --- | --- | --- | --- | --- |
| TBI at 3-months post injury, 0.01 | TBI at 6-months post injury, 0.01 | 0.051 | 4.733 | **<0.0001** |
|  | TBI at 12-months post injury, 0.01 | -0.008 | -0.719 | 1.000 |
| TBI at 3-months post injury, 0.03 | TBI at 6-months post injury, 0.03 | 0.027 | 2.482 | 1.000 |
|  | TBI at 12-months post injury, 0.03 | -0.005 | -0.447 | 1.000 |
| TBI at 3-months post injury, 0.06 | TBI at 6-months post injury, 0.06 | 0.009 | 0.830 | 1.000 |
|  | TBI at 12-months post injury, 0.06 | -0.003 | -0.311 | 1.000 |
| TBI at 3-months post injury, 0.09 | TBI at 6-months post injury, 0.09 | 0.003 | 0.262 | 1.000 |
|  | TBI at 12-months post injury, 0.09 | -0.003 | -0.256 | 1.000 |
| TBI at 3-months post injury, 0.12 | TBI at 6-months post injury, 0.12 | $6.805 \times 10^{-4}$ | 0.063 | 1.000 |
|  | TBI at 12-months post injury, 0.12 | -0.003 | -0.250 | 1.000 |
| TBI at 3-months post injury, 0.15 | TBI at 6-months post injury, 0.15 | $2.256 \times 10^{-4}$ | 0.021 | 1.000 |
|  | TBI at 12-months post injury, 0.15 | -0.002 | -0.219 | 1.000 |
| TBI at 3-months post injury, 0.18 | TBI at 6-months post injury, 0.18 | $1.249 \times 10^{-4}$ | 0.012 | 1.000 |
|  | TBI at 12-months post injury, 0.18 | -0.002 | -0.195 | 1.000 |
| TBI at 3-months post injury, 0.21 | TBI at 6-months post injury, 0.21 | $1.236 \times 10^{-4}$ | 0.011 | 1.000 |
|  | TBI at 12-months post injury, 0.21 | -0.002 | -0.173 | 1.000 |
| TBI at 3-months post injury, 0.24 | TBI at 6-months post injury, 0.24 | $2.759 \times 10^{-5}$ | 0.003 | 1.000 |
|  | TBI at 12-months post injury, 0.24 | -0.002 | -0.158 | 1.000 |

**Table S12.** Information Cascade in TBI patients over time. Results from the repeated measures ANCOVA with lesion volume as nuisance covariate (top table) and post hoc tests (bottom table).

| Effect | $F_{(2,20)}$ | p-val | Effect size ($\eta^2_{\text{partial}}$) |
| --- | --- | --- | --- |
| Time | 4.205 | 0.030 | 0.296 |

| Group 1 | Group 2 | Mean Difference | t | pval |
| --- | --- | --- | --- | --- |
| TBI at 3-months post-injury | TBI at 6-months post-injury | 0.010 | 2.195 | 0.080 |
|  | TBI at 12-months post-injury | -0.003 | -0.712 | 0.485 |
| TBI at 6-months post-injury | TBI at 12-months post-injury | -0.013 | -2.906 | **0.026** |

**Table S13.**
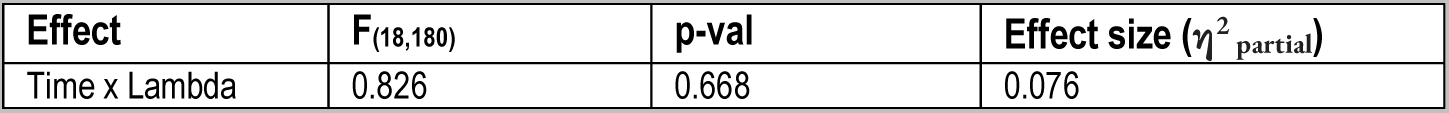
Information Transfer in TBI patients over time. Non-significant results from the repeated measures ANCOVA with lesion volume as nuisance covariate.

**Figure S1.**
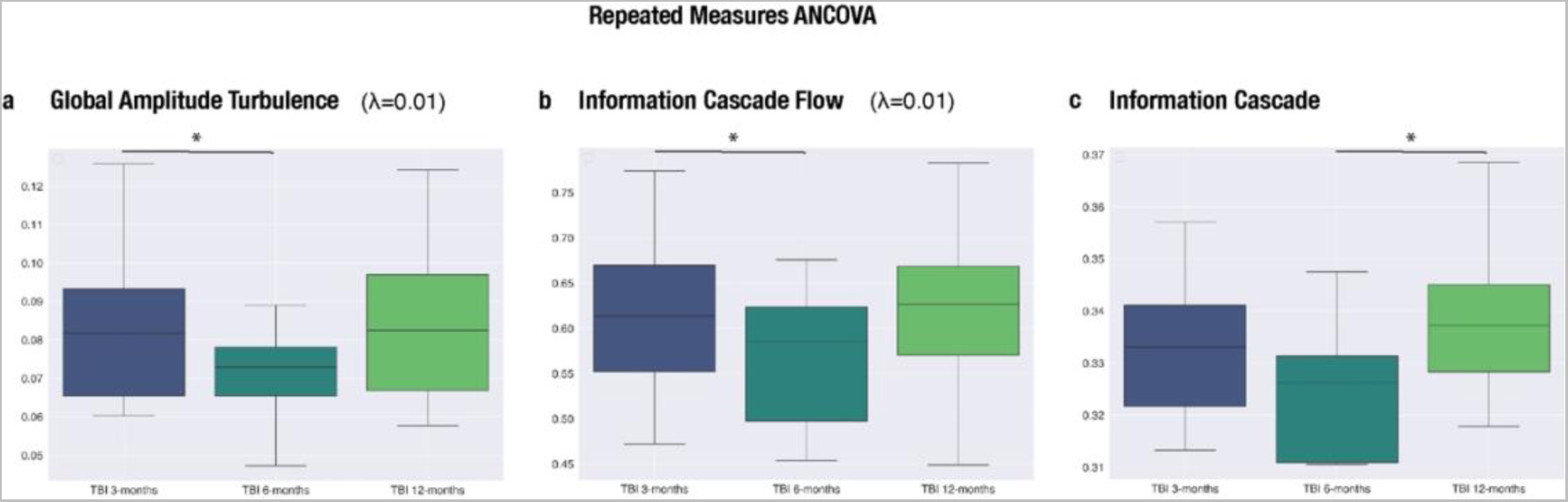
Boxplots depicting the significant results from the repeated measures ANCOVA with lesion volume as nuisance covariate.

**Tabla S14.** Amplitude turbulence in the 7 resting state networks (RSN) included in the Schaefer 1000 nodes parcellation. Results from the one-way ANOVA (4 groups) for λ=0.01 and λ=0.03 (top table) and post hoc tests for λ=0.03 (bottom table).

| Lambda | RSN | F<sub>(3,44)</sub> | p-val | Effect size ($\eta^2_{\text{partial}}$) |
| --- | --- | --- | --- | --- |
| 0.01 | VIS | 2.203 | 0.101 | 0.131 |
| 0.01 | SM | 2.014 | 0.126 | 0.121 |
| 0.01 | DAT | 2.211 | 0.100 | 0.131 |
| 0.01 | VAT | 1.820 | 0.157 | 0.110 |
| 0.01 | LIM | 1.486 | 0.231 | 0.092 |
| 0.01 | CNT | 1.931 | 0.139 | 0.116 |
| 0.01 | DMN | 1.881 | 0.147 | 0.114 |
| 0.03 | VIS | 2.859 | **0.048** | 0.163 |
| 0.03 | SM | 4.575 | **0.007** | 0.238 |
| 0.03 | DAT | 4.558 | **0.007** | 0.237 |
| 0.03 | VAT | 3.015 | **0.040** | 0.171 |
| 0.03 | LIM | 1.324 | 0.279 | 0.083 |
| 0.03 | CNT | 4.017 | **0.013** | 0.215 |
| 0.03 | DMN | 3.146 | **0.034** | 0.177 |

| RSN | Group 1 | Group 2 | Mean Difference | t | pval |
| --- | --- | --- | --- | --- | --- |
| VIS | Healthy Controls | TBI at 3-months post-injury | 0.015 | 2.244 | 0.127 |
|  |  | TBI at 6-months post-injury | 0.019 | 2.751 | **0.041** |
|  |  | TBI at 12-months post-injury | 0.012 | 1.713 | 0.329 |
| SM | Healthy Controls | TBI at 3-months post-injury | 0.011 | 1.983 | 0.210 |
|  |  | TBI at 6-months post-injury | 0.021 | 3.700 | **0.003** |
|  |  | TBI at 12-months post-injury | 0.010 | 1.795 | 0.289 |
| DAT | Healthy Controls | TBI at 3-months post-injury | 0.014 | 2.216 | 0.135 |
|  |  | TBI at 6-months post-injury | 0.023 | 3.660 | **0.004** |
|  |  | TBI at 12-months post-injury | 0.014 | 2.190 | 0.142 |
| VAT | Healthy Controls | TBI at 3-months post-injury | 0.007 | 1.361 | 0.530 |
|  |  | TBI at 6-months post-injury | 0.017 | 3.002 | **0.022** |
|  |  | TBI at 12-months post-injury | 0.008 | 1.373 | 0.523 |
| CNT | Healthy Controls | TBI at 3-months post-injury | 0.010 | 1.875 | 0.253 |
|  |  | TBI at 6-months post-injury | 0.019 | 3.466 | **0.006** |
|  |  | TBI at 12-months post-injury | 0.010 | 1.865 | 0.258 |
| DMN | Healthy Controls | TBI at 3-months post-injury | 0.010 | 1.682 | 0.345 |
|  |  | TBI at 6-months post-injury | 0.018 | 3.067 | **0.019** |
|  |  | TBI at 12-months post-injury | 0.009 | 1.554 | 0.415 |

**Table S15.**
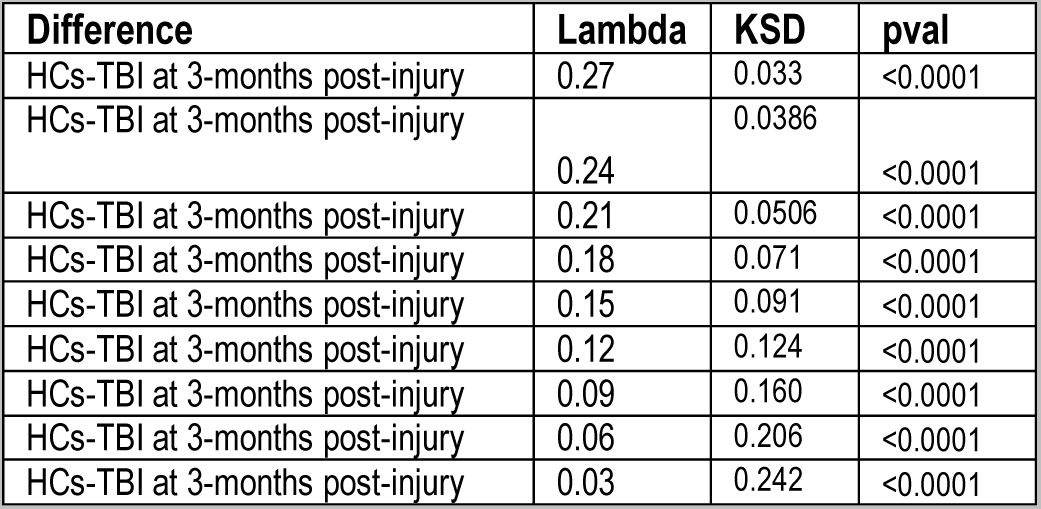
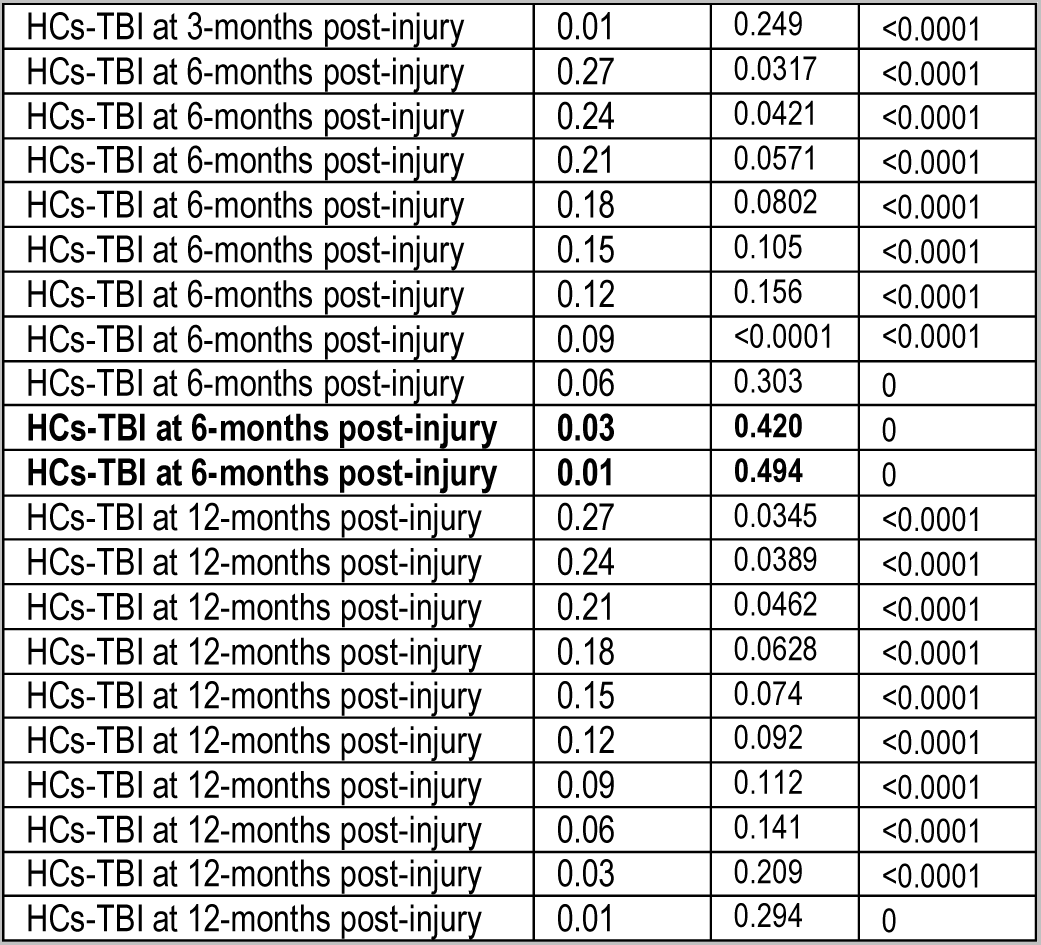
Node-level amplitude turbulence. Results from the Kolmogorov-Smirnov test.

Table S16. Correlation between turbulent-like dynamics measures and cognitive performance in TBI patients at 3-months post-injury. The variables r and p denote the Pearson correlation coefficient and associated p-value, respectively. The asterisk (*) denotes that the correlation survived FDR correction for multiple comparisons across tasks (p-adj < 0.05). C: Color, CW: Color-Word, TMTA: Trail Making Test A, TMTB: Trail Making Test B, Dig: Digit Span Forward and Backward, VSAT: Visual Search and Attention Test.

**Table S16. Correlation between turbulent-like dynamics measures and cognitive performance in TBI patients at 3-months post-injury.**
| Turbulent measure | Stroop C total |  | Stroop CW total |  | TMTA |  | TMTB |  | Dig Total |  | VSAT total |  |
| --- | --- | --- | --- | --- | --- | --- | --- | --- | --- | --- | --- | --- |
|  | r | p | r | p | r | p | r | p | r | p | r | p |
| Turbulence Global $\lambda=0.06$ | 0.372 | 0.260 | 0.623 | 0.054 | 0.254 | 0.451 | 0.198 | 0.583 | 0.679 | 0.022 | 0.275 | 0.412 |
| Turbulence Global $\lambda=0.03$ | 0.416 | 0.203 | 0.548 | 0.101 | 0.235 | 0.486 | 0.267 | 0.456 | 0.711 | 0.014 | 0.163 | 0.633 |
| Turbulence Global $\lambda=0.01$ | 0.442 | 0.173 | 0.443 | 0.200 | 0.181 | 0.595 | 0.331 | 0.351 | 0.685 | 0.020 | 0.093 | 0.785 |
| Info Casc Flow $\lambda=0.01$ | 0.584 | 0.059 | 0.346 | 0.327 | 0.159 | 0.641 | 0.206 | 0.569 | 0.754 | **0.007\*** | -0.047 | 0.891 |
| Info Transfer $\lambda=0.27$ | -0.059 | 0.864 | 0.120 | 0.741 | 0.360 | 0.276 | 0.324 | 0.360 | -0.023 | 0.945 | 0.196 | 0.563 |
| Info Transfer $\lambda=0.24$ | -0.034 | 0.921 | 0.128 | 0.725 | 0.433 | 0.184 | 0.391 | 0.264 | -0.038 | 0.911 | 0.217 | 0.521 |
| Info Transfer $\lambda=0.21$ | -0.047 | 0.891 | 0.101 | 0.780 | 0.424 | 0.194 | 0.372 | 0.290 | -0.004 | 0.991 | 0.196 | 0.563 |
| Info Transfer $\lambda=0.18$ | -0.059 | 0.863 | 0.062 | 0.865 | 0.415 | 0.204 | 0.343 | 0.332 | 0.014 | 0.967 | 0.177 | 0.602 |
| Info Transfer $\lambda=0.15$ | -0.094 | 0.784 | 0.011 | 0.976 | 0.415 | 0.204 | 0.276 | 0.441 | -0.007 | 0.985 | 0.158 | 0.642 |

**Table S17.** Correlation between turbulent-like dynamics measures by resting-state network (RSN) and cognitive performance in TBI patients at 3-months post-injury. The variables r and p denote the Pearson correlation coefficient and associated p-value, respectively. The asterisk (*) denotes that the correlation survived FDR correction for multiple comparisons across tasks (p-adj < 0.05). C: Color, CW: Color-Word, TMTA: Trail Making Test A, TMTB: Trail Making Test B, Dig: Digit Span Forward and Backward, VSAT: Visual Search and Attention Test.

| Turbulent measure by RSN | Stroop C total |  | Stroop CW total |  | TMTA |  | TMTB |  | Dig Total |  | VSAT total |  |
| --- | --- | --- | --- | --- | --- | --- | --- | --- | --- | --- | --- | --- |
|  | r | p | r | p | r | p | r | p | r | p | r | p |
| Turbulence VIS $\lambda=0.03$ | 0.526 | 0.096 | 0.728 | 0.017 | 0.153 | 0.653 | 0.157 | 0.665 | 0.689 | 0.019 | 0.260 | 0.440 |
| Turbulence SM $\lambda=0.03$ | 0.375 | 0.413 | 0.392 | 0.263 | 0.306 | 0.360 | 0.395 | 0.259 | 0.679 | 0.022 | 0.147 | 0.665 |
| Turbulence DAT $\lambda=0.03$ | 0.256 | 0.448 | 0.545 | 0.103 | 0.273 | 0.417 | 0.267 | 0.456 | 0.614 | 0.044 | 0.211 | 0.534 |
| Turbulence VAT $\lambda=0.03$ | 0.416 | 0.203 | 0.436 | 0.208 | 0.275 | 0.413 | 0.346 | 0.328 | 0.721 | 0.012 | 0.111 | 0.746 |
| Turbulence LIM $\lambda=0.03$ | 0.521 | 0.100 | 0.412 | 0.236 | 0.136 | 0.691 | 0.114 | 0.754 | 0.675 | 0.023 | 0.046 | 0.893 |
| Turbulence CON $\lambda=0.03$ | 0.374 | 0.257 | 0.496 | 0.145 | 0.218 | 0.520 | 0.272 | 0.447 | 0.633 | 0.036 | 0.178 | 0.600 |
| Turbulence DMN $\lambda=0.03$ | 0.495 | 0.121 | 0.491 | 0.150 | 0.163 | 0.632 | 0.163 | 0.653 | 0.744 | **0.009\*** | 0.081 | 0.813 |

**Table S18.**
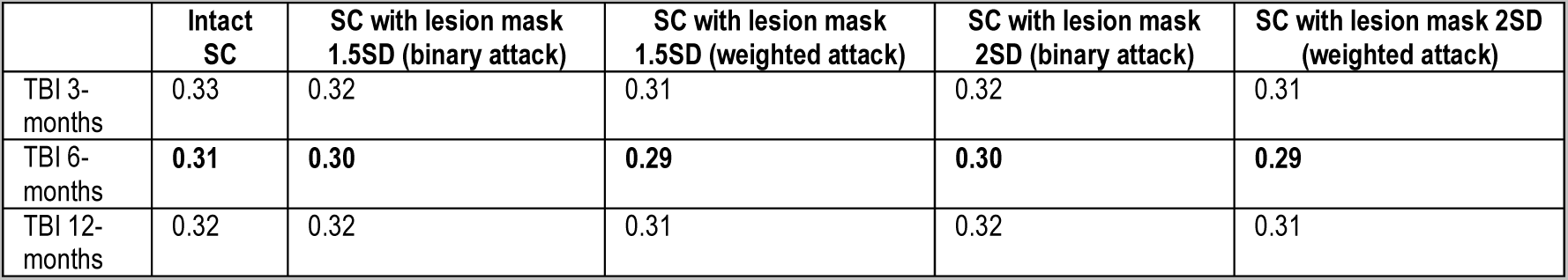
Global coupling parameter after the simulated attack approach. SC: structural connectivity (see Material and Methods).

|  | Intact SC | SC with lesion mask 1.5SD (binary attack) | SC with lesion mask 1.5SD (weighted attack) | SC with lesion mask 2SD (binary attack) | SC with lesion mask 2SD (weighted attack) |
| --- | --- | --- | --- | --- | --- |
| TBI 3-months | 0.33 | 0.32 | 0.31 | 0.32 | 0.31 |
| TBI 6-months | **0.31** | **0.30** | **0.29** | **0.30** | **0.29** |
| TBI 12-months | 0.32 | 0.32 | 0.31 | 0.32 | 0.31 |

**Table S19.** Anatomical regions lesions in the structural connectivity matrix thresholded at 1.5SD. The node label is indicated along with the corresponding anatomical region (see Material and Methods).

| Node to attack Schaefer 1000 nodes label | Anatomical region | Frequency (TBI Patients with lesion) |
| --- | --- | --- |
| 143 | 7Networks_LH_SomMot_62\ | 1 |
| 155 | 7Networks_LH_SomMot_74\ | 1 |
| 161 | 7Networks_LH_SomMot_80\ | 1 |
| 189 | 7Networks_LH_DorsAttn_Post_17\ | 1 |
| 245 | 7Networks_LH_SalVentAttn_FrOperIns_1\ | 1 |
| 249 | 7Networks_LH_SalVentAttn_FrOperIns_5\ | 1 |
| 259 | 7Networks_LH_SalVentAttn_FrOperIns_15\ | 1 |
| 274 | 7Networks_LH_SalVentAttn_Med_5\ | 1 |
| 284 | 7Networks_LH_SalVentAttn_Med_15\ | 1 |
| 292 | 7Networks_LH_Limbic_OFC_4\ | 1 |
| 367 | 7Networks_LH_Cont_Cing_3\ | 1 |
| 368 | 7Networks_LH_Cont_Cing_4\ | 1 |
| 370 | 7Networks_LH_Cont_Cing_6\ | 1 |
| 416 | 7Networks_LH_Default_PFC_1\ | 2 |
| 417 | 7Networks_LH_Default_PFC_2\ | 1 |
| 471 | 7Networks_LH_Default_pCunPCC_6\ | 1 |
| 497 | 7Networks_LH_Default_pCunPCC_32\ | 1 |
| 533 | 7Networks_RH_Vis_33\ | 1 |
| 536 | 7Networks_RH_Vis_36\ | 1 |
| 555 | 7Networks_RH_Vis_55\ | 1 |
| 586 | 7Networks_RH_SomMot_5\ | 1 |
| 591 | 7Networks_RH_SomMot_10\ | 1 |
| 598 | 7Networks_RH_SomMot_17\ | 1 |
| 612 | 7Networks_RH_SomMot_31\ | 1 |
| 618 | 7Networks_RH_SomMot_37\ | 1 |
| 632 | 7Networks_RH_SomMot_51\ | 1 |
| 671 | 7Networks_RH_SomMot_90\ | 1 |
| 679 | 7Networks_RH_SomMot_98\ | 1 |
| 717 | 7Networks_RH_DorsAttn_Post_33\ | 1 |
| 722 | 7Networks_RH_DorsAttn_Post_38\ | 1 |
| 753 | 7Networks_RH_SalVentAttn_TempOccPar_8\ | 1 |
| 775 | 7Networks_RH_SalVentAttn_FrOperIns_10\ | 1 |
| 805 | 7Networks_RH_SalVentAttn_Med_14\ | 1 |
| 842 | 7Networks_RH_Limbic_TempPole_17\ | 1 |
| 843 | 7Networks_RH_Cont_Par_1\ | 1 |
| 865 | 7Networks_RH_Cont_PFCv_2\ | 2 |
| 913 | 7Networks_RH_Default_Par_1\ | 1 |
| 919 | 7Networks_RH_Default_Par_7\ | 1 |
| 949 | 7Networks_RH_Default_PFCv_3\ | 1 |
| 961 | 7Networks_RH_Default_PFCdPFCm_5\ | 1 |
| 999 | 7Networks_RH_Cont_pCun_2\ | 1 |

**Figure S2.**
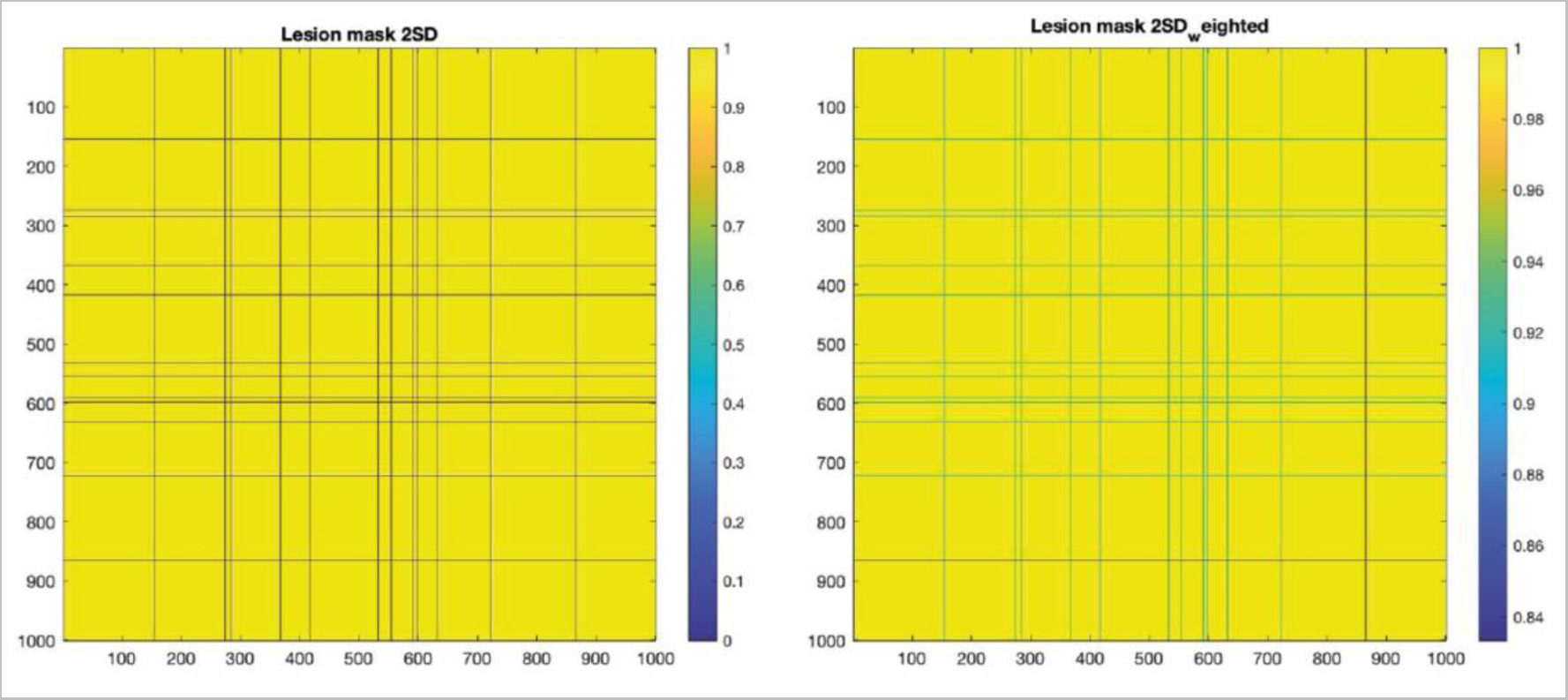
Heatmaps for the lesion mask thresholded at 2SD with the binary (left) and weighted (right) approach for the simulated attack (see Material and Methods).

**Table S20.** Susceptibility in HCs (session 1 and 2) vs TBI (3-,6- and 12-months post-injury). Results from the two-sided Wilcoxon rank sum test. Lesion mask 1.5 SD (see Material and Methods).

| Group comparison | Z | pval | padj |
| --- | --- | --- | --- |
| HCs s1 vs TBI 3mo | 6.29 | <0.0001 | <0.0001 |
| HCs s1 vs TBI 6mo | 10.77 | <0.0001 | <0.0001 |
| HCs s1 vs TBI 12mo | 10.25 | <0.0001 | <0.0001 |
| HCs s1 vs TBI 3mo lesion 1.5 SD Binary | 10.25 | <0.0001 | <0.0001 |
| HCs s1 vs TBI 6mo lesion 1.5 SD Binary | 11.89 | <0.0001 | <0.0001 |
| HCs s1 vs TBI 12mo lesion 1.5 SD Binary | 11.58 | <0.0001 | <0.0001 |
| HCs s1 vs TBI 3mo lesion 1.5 SD Weighted | 7.09 | <0.0001 | <0.0001 |
| HCs s1 vs TBI 6mo lesion 1.5 SD Weighted | 11.09 | <0.0001 | <0.0001 |
| HCs s1 vs TBI 12mo lesion 1.5 SD Weighted | 10 | <0.0001 | <0.0001 |
| HCs s2 vs TBI 3mo | 2.80 | 0.0051 | 0.0051 |
| HCs s2 vs TBI 6mo | 9.04 | <0.0001 | <0.0001 |
| HCs s2 vs TBI 12mo | 8.03 | <0.0001 | <0.0001 |
| HCs s2 vs TBI 3mo lesion 1.5 SD Binary | 8.21 | <0.0001 | <0.0001 |
| HCs s2 vs TBI 6mo lesion 1.5 SD Binary | 11.44 | <0.0001 | <0.0001 |
| HCs s2 vs TBI 12mo lesion 1.5 SD Binary | 10.71 | <0.0001 | <0.0001 |
| HCs s2 vs TBI 3mo lesion 1.5 SD Weighted | 3.83 | <0.0001 | <0.0001 |
| HCs s2 vs TBI 6mo lesion 1.5 SD Weighted | 9.68 | <0.0001 | <0.0001 |
| HCs s2 vs TBI 12mo lesion 1.5 SD Weighted | 7.66 | <0.0001 | <0.0001 |

**Table S21.** Information encoding capability in HCs (session 1 and 2) vs TBI (3-,6- and 12-months post-injury). Results from the two-sided Wilcoxon rank sum test. Lesion mask 1.5 SD (see Material and Methods).

| Group comparison | Z | pval | padj |
| --- | --- | --- | --- |
| HCs s1 vs TBI 3mo | 8.35 | <0.0001 | <0.0001 |
| HCs s1 vs TBI 6mo | 12.02 | <0.0001 | <0.0001 |
| HCs s1 vs TBI 12mo | 11.82 | <0.0001 | <0.0001 |
| HCs s1 vs TBI 3mo lesion 1.5 SD Binary | 10.72 | <0.0001 | <0.0001 |
| HCs s1 vs TBI 6mo lesion 1.5 SD Binary | 12.18 | <0.0001 | <0.0001 |
| HCs s1 vs TBI 12mo lesion 1.5 SD Binary | 12.07 | <0.0001 | <0.0001 |
| HCs s1 vs TBI 3mo lesion 1.5 SD Weighted | 9.25 | <0.0001 | <0.0001 |
| HCs s1 vs TBI 6mo lesion 1.5 SD Weighted | 12.10 | <0.0001 | <0.0001 |
| HCs s1 vs TBI 12mo lesion 1.5 SD Weighted | 11.71 | <0.0001 | <0.0001 |
| HCs s2 vs TBI 3mo | 4.59 | <0.0001 | <0.0001 |
| HCs s2 vs TBI 6mo | 11.36 | <0.0001 | <0.0001 |
| HCs s2 vs TBI 12mo | 10.80 | <0.0001 | <0.0001 |
| HCs s2 vs TBI 3mo lesion 1.5 SD Binary | 8.64 | <0.0001 | <0.0001 |
| HCs s2 vs TBI 6mo lesion 1.5 SD Binary | 12.04 | <0.0001 | <0.0001 |
| HCs s2 vs TBI 12mo lesion 1.5 SD Binary | 11.60 | <0.0001 | <0.0001 |
| HCs s2 vs TBI 3mo lesion 1.5 SD Weighted | 5.86 | <0.0001 | <0.0001 |
| HCs s2 vs TBI 6mo lesion 1.5 SD Weighted | 11.73 | <0.0001 | <0.0001 |
| HCs s2 vs TBI 12mo lesion 1.5 SD Weighted | 10.49 | <0.0001 | <0.0001 |

**Table S22.** Susceptibility in HCs (session 1 and 2) vs TBI (3-,6- and 12-months post-injury). Results from the two-sided Wilcoxon rank sum test. Lesion mask 2 SD (see Material and Methods).

| Group comparison | Z | pval | padj |
| --- | --- | --- | --- |
| HCs s1 vs TBI 3mo | 6.29 | <0.0001 | <0.0001 |
| HCs s1 vs TBI 6mo | 10.77 | <0.0001 | <0.0001 |
| HCs s1 vs TBI 12mo | 10.25 | <0.0001 | <0.0001 |
| HCs s1 vs TBI 3mo lesion 2 SD Binary | 8.54 | <0.0001 | <0.0001 |
| HCs s1 vs TBI 6mo lesion 2 SD Binary | 11.46 | <0.0001 | <0.0001 |
| HCs s1 vs TBI 12mo lesion 2 SD Binary | 10.80 | <0.0001 | <0.0001 |
| HCs s1 vs TBI 3mo lesion 2 SD Weighted | 6.93 | <0.0001 | <0.0001 |
| HCs s1 vs TBI 6mo lesion 2 SD Weighted | 11.06 | <0.0001 | <0.0001 |
| HCs s1 vs TBI 12mo lesion 2 SD Weighted | 9.90 | <0.0001 | <0.0001 |
| HCs s2 vs TBI 3mo | 2.80 | 0.0051 | 0.0051 |
| HCs s2 vs TBI 6mo | 9.04 | <0.0001 | <0.0001 |
| HCs s2 vs TBI 12mo | 8.03 | <0.0001 | <0.0001 |
| HCs s2 vs TBI 3mo lesion 2 SD Binary | 5.76 | <0.0001 | <0.0001 |
| HCs s2 vs TBI 6mo lesion 2 SD Binary | 10.49 | <0.0001 | <0.0001 |
| HCs s2 vs TBI 12mo lesion 2 SD Binary | 9.17 | <0.0001 | <0.0001 |
| HCs s2 vs TBI 3mo lesion 2 SD Weighted | 3.63 | 0.0002 | <0.0001 |
| HCs s2 vs TBI 6mo lesion 2 SD Weighted | 9.61 | <0.0001 | <0.0001 |
| HCs s2 vs TBI 12mo lesion 2 SD Weighted | 7.49 | <0.0001 | <0.0001 |

**Table S23.**
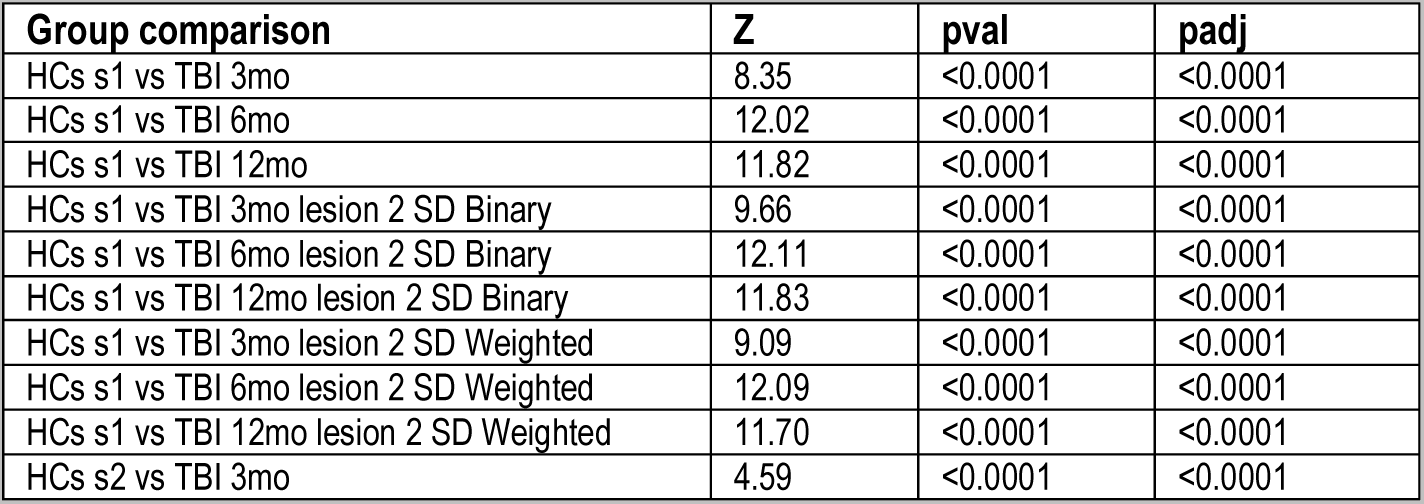
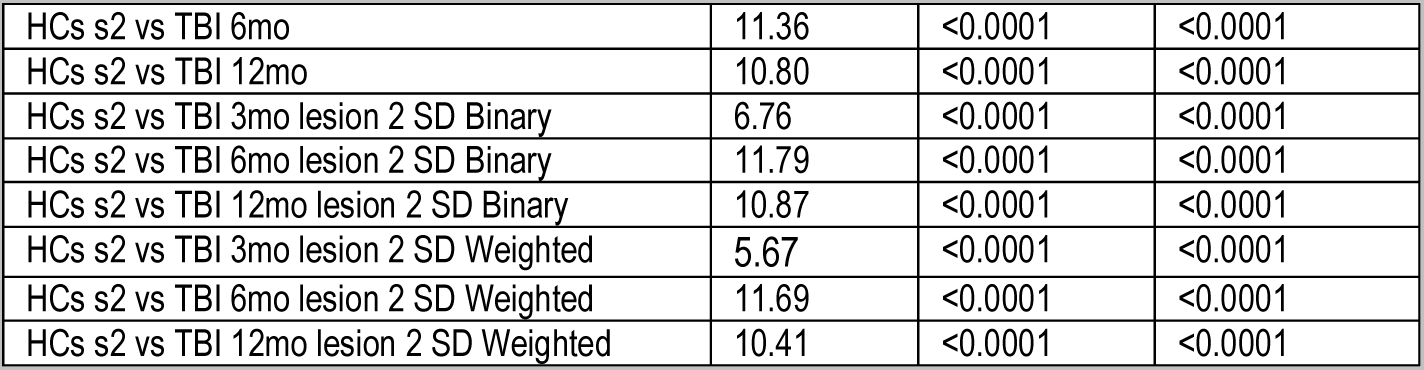
Information encoding capability in HCs (session 1 and 2) vs TBI (3-,6- and 12-months post-injury). Results from the two-sided Wilcoxon rank sum test. Lesion mask 2 SD (see Material and Methods).

| Group comparison | Z | pval | padj |
| --- | --- | --- | --- |
| HCs s1 vs TBI 3mo | 8.35 | <0.0001 | <0.0001 |
| HCs s1 vs TBI 6mo | 12.02 | <0.0001 | <0.0001 |
| HCs s1 vs TBI 12mo | 11.82 | <0.0001 | <0.0001 |
| HCs s1 vs TBI 3mo lesion 2 SD Binary | 9.66 | <0.0001 | <0.0001 |
| HCs s1 vs TBI 6mo lesion 2 SD Binary | 12.11 | <0.0001 | <0.0001 |
| HCs s1 vs TBI 12mo lesion 2 SD Binary | 11.83 | <0.0001 | <0.0001 |
| HCs s1 vs TBI 3mo lesion 2 SD Weighted | 9.09 | <0.0001 | <0.0001 |
| HCs s1 vs TBI 6mo lesion 2 SD Weighted | 12.09 | <0.0001 | <0.0001 |
| HCs s1 vs TBI 12mo lesion 2 SD Weighted | 11.70 | <0.0001 | <0.0001 |
| HCs s2 vs TBI 3mo | 4.59 | <0.0001 | <0.0001 |
| HCS s2 vs TBI 6mo | 11.36 | <0.0001 | <0.0001 |
| HCS s2 vs TBI 12mo | 10.80 | <0.0001 | <0.0001 |
| HCS s2 vs TBI 3mo lesion 2 SD Binary | 6.76 | <0.0001 | <0.0001 |
| HCS s2 vs TBI 6mo lesion 2 SD Binary | 11.79 | <0.0001 | <0.0001 |
| HCS s2 vs TBI 12mo lesion 2 SD Binary | 10.87 | <0.0001 | <0.0001 |
| HCS s2 vs TBI 3mo lesion 2 SD Weighted | 5.67 | <0.0001 | <0.0001 |
| HCS s2 vs TBI 6mo lesion 2 SD Weighted | 11.69 | <0.0001 | <0.0001 |
| HCS s2 vs TBI 12mo lesion 2 SD Weighted | 10.41 | <0.0001 | <0.0001 |

